# Beyond natural amino acids: Extending immunogenicity risk assessment to non-canonical peptide drugs through chemical feature encoding

**DOI:** 10.64898/2026.05.22.727138

**Authors:** Maria Cairoli, Morten Nielsen, Catherine Betts, Olga Obrezanova, Leonardo De Maria

**Affiliations:** Discovery Computation and AI, Hit Discovery, Discovery Sciences, Biopharmaceuticals R&D, AstraZeneca, Gothenburg, Sweden; Department of Health Technology, Technical University of Denmark, Lyngby, Denmark; Immune Safety, Clinical Pharmacology and Safety Sciences, Biopharmaceuticals R&D, AstraZeneca, Cambridge, UK; Biologics Engineering, Oncology R&D, AstraZeneca, Cambridge, UK

**Keywords:** immunogenicity prediction, MHC class II, non-natural amino acids, post-translational modifications, machine learning, chemical fingerprints

## Abstract

Peptide therapeutics are increasingly used to treat challenging diseases, but immunogenicity risks limit their clinical success. *In silico* tools enable immunogenicity screening through prediction of peptide-MHCII binding, yet current methods fail to capture chemical properties of non-natural amino acids routinely incorporated to improve drug properties. Here, we present a machine learning approach combining chemical fingerprints with sequence information to predict MHC class II binding for both canonical and modified peptides. We propose two molecular representations (direct-encoding and similarity-based chemical fingerprints) that preserve positional information while encoding chemical diversity. These representations achieved performance comparable to sequence-based encodings (BLOSUM62 and one-hot) for canonical peptides while accurately identifying binding cores and motifs. Testing on citrullinated peptides, chemical fingerprints substantially improved quantitative prediction accuracy while maintaining comparable linear correlation across encoding methods, demonstrating the importance of explicit chemical representation for accurate absolute binding affinity prediction. These descriptors can be integrated into pan-allele prediction frameworks, enabling immunogenicity risk assessment across diverse modifications and therapeutic modalities, including peptide therapeutics, antibody-drug conjugates, and synthetic vaccines. The proposed chemistry-informed framework addresses a critical gap in preclinical drug development, facilitating early mitigation strategies before costly clinical trials.

## 1. Introduction

Peptide therapeutics are increasingly relevant in the global pharmaceutical industry, playing a key role in treating some of the most challenging diseases, including diabetes, cancer and multiple sclerosis (1–3). Their intermediate size between small molecules and biologics confers advantageous properties such as high target specificity and minimal off-target effects (4). However, the clinical efficacy of such therapeutics is often limited by suboptimal properties, including short in vivo half-lives, poor oral bioavailability, limited permeation across biological barriers, and immunogenicity (2, 5). To improve these properties while preserving biological activity, medicinal chemists increasingly incorporate non-natural amino acids (NNAAs) beyond the 20 canonical building blocks (4, 6, 7). However, despite these advances, the extent to which non-natural amino acids affect immunogenicity remains largely unknown (8, 9), representing a critical concern for the success of clinical trials (10).

Immunogenicity refers to an unwanted immune response to protein or peptide biologics (11). Such response occurs when drug peptides are presented on major histocompatibility complex (MHC) molecules to T cells, leading to their activation, proliferation and differentiation into effector cells that can recognize and eliminate the antigen (12). In this process, peptide-MHC (pMHC) binding is the key factor determining which peptides will ultimately be presented to T cells and trigger an immune response (12). Predicting the binding to MHC class II molecules helps identify potential T cell epitopes that can drive the critical T-cell dependent B cell response, which is crucial for anti-drug antibody (ADA) production.

*In silico* methods have been developed to predict pMHCII binding and recognize potentially immunogenic peptides (13–15), and have been demonstrated powerful for preclinical immunogenicity assessment before costly clinical trials (16). These tools simultaneously address peptide alignment and binding motif characterization due to the open-ended MHC class II binding groove, which accommodates variable-length peptides and allows flanking regions to influence binding affinity (12). The introduction of NNAAs could affect these binding interactions and potentially trigger T cell activation (8, 10). However, current prediction algorithms are primarily trained on the 20 natural amino acids, and may not accurately predict binding for NNAA-based peptides (10). Despite models encoding unnatural amino acids have been proposed (10, 17), they fail to capture the chemical diversity of the modified residues. For instance, Alvarez et al. used sparse encoding to model phosphorylated peptides in NetMHCIIphosPan (17), while Mattei et al. encoded unnatural amino acids with a neutral place holder ‘X’, supporting only one modification type (10). Excluding explicit chemical information limits the ability to distinguish among chemically diverse non-natural residues, potentially hampering accurate binding affinity prediction and preventing the inclusion of a broader range of synthetic modifications that medicinal chemists routinely employ in modern peptide therapeutic design.

Chemical fingerprints such as Extended Connectivity Fingerprints (ECFP) (18) and MinHashed Atom-Pair Chiral (MAPC) (19) are increasingly used in machine learning applications to describe peptides (20–22). These are derived from Simplified Molecular Input Line Entry System (SMILES) representation, which enables the description of molecular structures for peptides including both natural and non-natural residues. To the best of our knowledge, however, these fingerprints have not yet been tested on immunogenicity prediction. Unlike sequence-based representations, canonical fingerprint applications encode entire peptides as unified molecular entities, which may not explicitly preserve the positional arrangement of amino acids critical for binding core identification.

Here, we evaluate the effectiveness of chemical fingerprints as alternative molecular representations for *in silico* pMHCII prediction against sequence-based encodings. We compare fingerprints that encode peptides as single molecular entities and propose two novel representations that combine chemical diversity with positional information. We focus on single-allele models to isolate the effects of molecular representation from pan-allele learning. We benchmark our approaches against established sequence-based encodings, validate performance on natural amino acids across MHC class II and I alleles, and ultimately assess performance on available NNAA-based peptides, which in this study include citrullinated peptides. The proposed representations operate on the same principle as substitution matrices, enabling seamless integration into pan-allele prediction methods such as NetMHCIIpan. By addressing a critical gap in immunoinformatics, this work establishes a flexible, chemistry-informed framework for immunogenicity prediction across the diverse chemical space of modern peptide therapeutics.

## 2. Methods

### 2.1 Data sets

#### Peptide-MHC class II binding affinity data sets

We retrieved binding affinity data for peptide sequences from the Immune Epitope Database (IEDB, www.iedb.org) (23). For peptides including natural amino acids, we employed the data set curated by Nielsen et al. (14), encompassing binding affinity data for several alleles (**Table 1**). We retrieved NNAA-based peptides from the IEDB database, retaining those having quantitative binding affinity data (IC50) and clearly defined modification. This allowed retaining 225 citrullinated peptides for DRB1*01:01 and 283 citrullinated peptides for DRB1*04:01. We merged modified peptides with NA-based peptides following the ‘Hobohm1’ (24) inspired algorithm proposed by Nielsen et al. (13) to minimize the overlap between training and evaluation data. This algorithm creates non-redundant peptide clusters by grouping sequences that share identical 9-mer overlaps, then splits these clusters into five cross-validation subsets to minimize sequence redundancy between training and test data (13). We tokenized peptide sequences at the residue level, with canonical amino acids represented by single-letter codes and non-canonical amino acids enclosed in square brackets (e.g., [Cit] for citrulline), using a pattern-matching algorithm to identify and extract tokens from peptide sequences. We applied this tokenization scheme consistently across all algorithms.

**Table 1.**
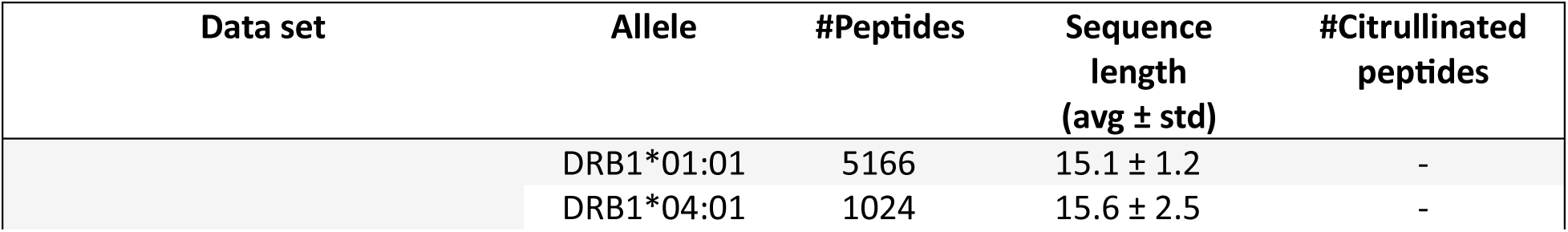
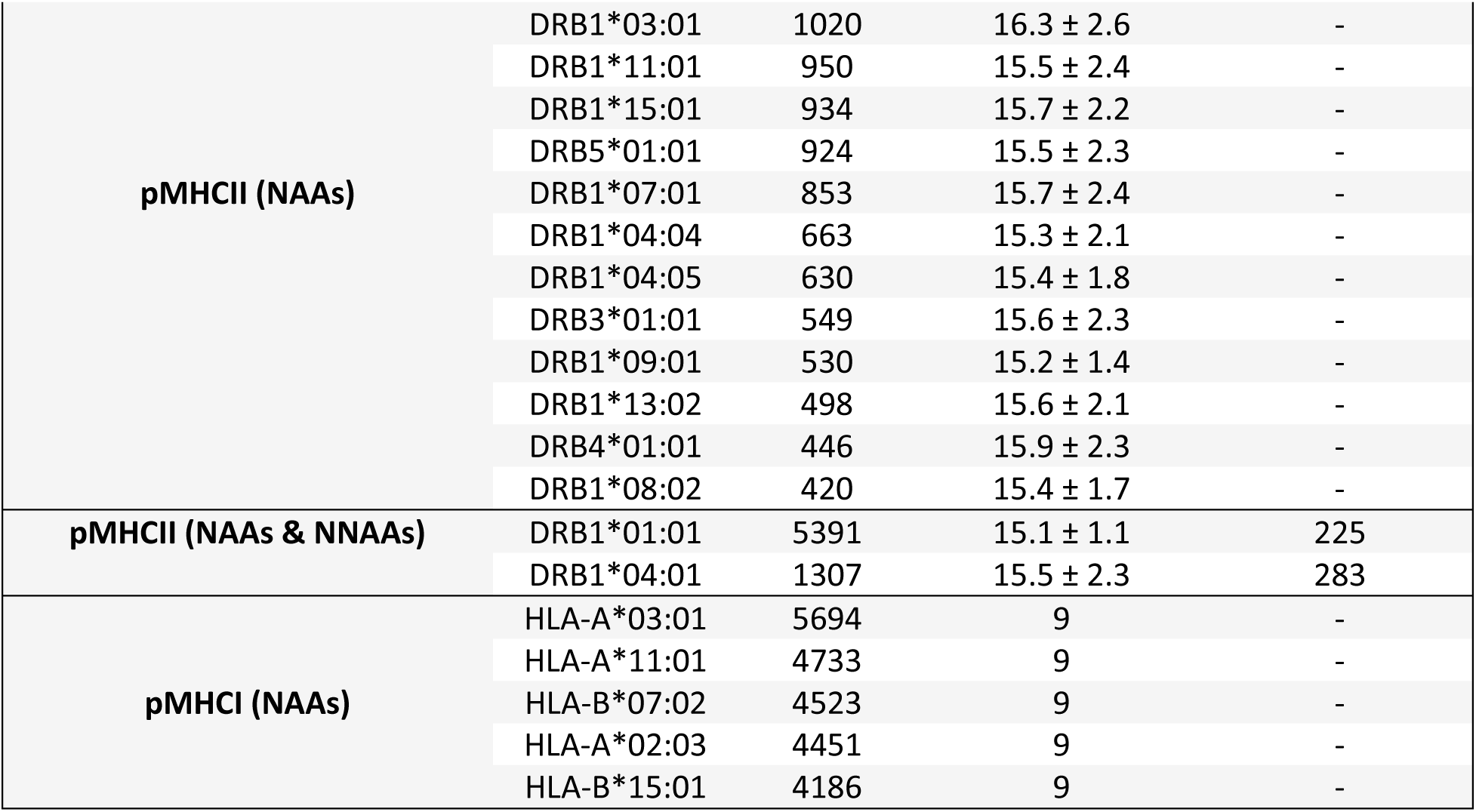
Details of training data sets. pMHCII, pMHCI: peptide-MHC class II and I binding; NAAs: natural amino acids; NNAAs: non-natural amino acids.

The binding affinity score was computed as

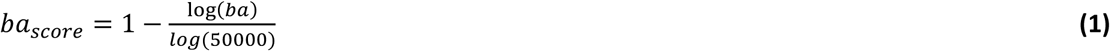

where ba is the binding affinity (IC50) measured in nM (14). ba_score_ values lower than 0 and higher than 1 were clipped to [0,1] range to avoid bias. **Table 1** reports details on these data sets.

#### Peptide-MHC class I binding affinity data set

We considered the dataset curated by Reynisson et al. (15) to retrieve quantitative binding affinity (BA) data for MHC class I alleles. We retained only peptides with a length equal to 9 amino acids, for five alleles encompassing both A and B allotypes: HLA-A*03:01, HLA-A*11:01, HLA-B*07:02, HLA-A*02:03, HLA-B*15:01 (**Table 1**). Data were clustered in five folds before prediction using the ‘Hobohm1’ inspired algorithm described above.

#### Data set from Human Proteome

Predicting random sequences can be useful to determining a baseline or background distribution against which the conservation of the amino acid patterns in immunogenic peptides can be established. For NAA-based peptides, we extracted 20,420 reviewed (Swiss-Prot) sequences from the UniProt Knowledgebase (UniProtKB; https://www.uniprot.org/) (25), considering ‘Human’ as organism. We selected sequences that only include the 20 natural amino acids, which were then split into sequences of 9-mers and 15-mers for MHC class I and class II alleles, obtaining 10,398,028 and 10,584,973 sequences, respectively. For each MHC class, we randomly sampled 500,000 sequences ultimately employed for prediction.

#### Data set from Rebak et al

We extracted modified sequences from the dataset published by Rebak et al. (26) for sequence logo estimation of modified residues, as it encompasses 14,056 citrullination sites within 4,008 proteins (26). We selected only sequences containing citrullinated sites, excluding those including other modifications. Sequences longer than 15 amino acids were split in 15-mers. We excluded duplicates, sequences shorter than 9 amino acids and not containing any modification, obtaining a total of 43,894 sequences considered for prediction.

### 2.2 Molecular representations

We evaluated three chemical fingerprint-based approaches to encode peptides: peptide-level, direct-encoding, and similarity-based fingerprints. These were benchmarked against established sequence-based encoding methods for binding affinity prediction and binding motif recognition.

#### Sequence-based encodings

##### BLOSUM Matrix

BLOSUM (block substitution matrix) (27) matrices are the state-of-the-art in immunoinformatics to encode amino acid sequences. These matrices are based on local alignments of conserved protein regions and are built by calculating the log-odds scores for amino acid substitutions observed in blocks of aligned sequences from the Blocks Database (28). Each BLOSUM matrix is derived from sequence alignments clustered at different percent identity thresholds, for the 20 natural amino acids. In this study we consider BLOSUM62, which uses sequences with no more than 62% identity.

##### One-hot encoding

This method represents each amino acid as a unique binary vector of length 20 (for natural amino acids), where a single position is set to 1 and all other positions are 0. This encoding captures only categorical information without incorporating physicochemical properties or evolutionary relationships.

##### NNAAIndex

The Index of Natural and Non-Natural Amino Acids (NNAAIndex) (29) is a structural matrix that characterizes 155 properties of 615 amino acids, including both natural and non-natural amino acids. These properties were clustered into 6 representative patterns that encompass geometric characteristics for the considered amino acids, H-bond, connectivity, accessible surface area, integy moments index, volume and shape (29). While this descriptor encodes chemical information, it is limited to a fixed vocabulary of pre-characterized amino acids and uses sequence information. We therefore categorize it as a sequence-based encoding in this manuscript.

#### Peptide-level fingerprints

We evaluated 10 peptide-level chemical fingerprints (**Table 2**) representing different encoding classes (30), each capturing diverse chemical properties. These fingerprints encode the entire peptide as a single molecular entity.

**Table 2.**
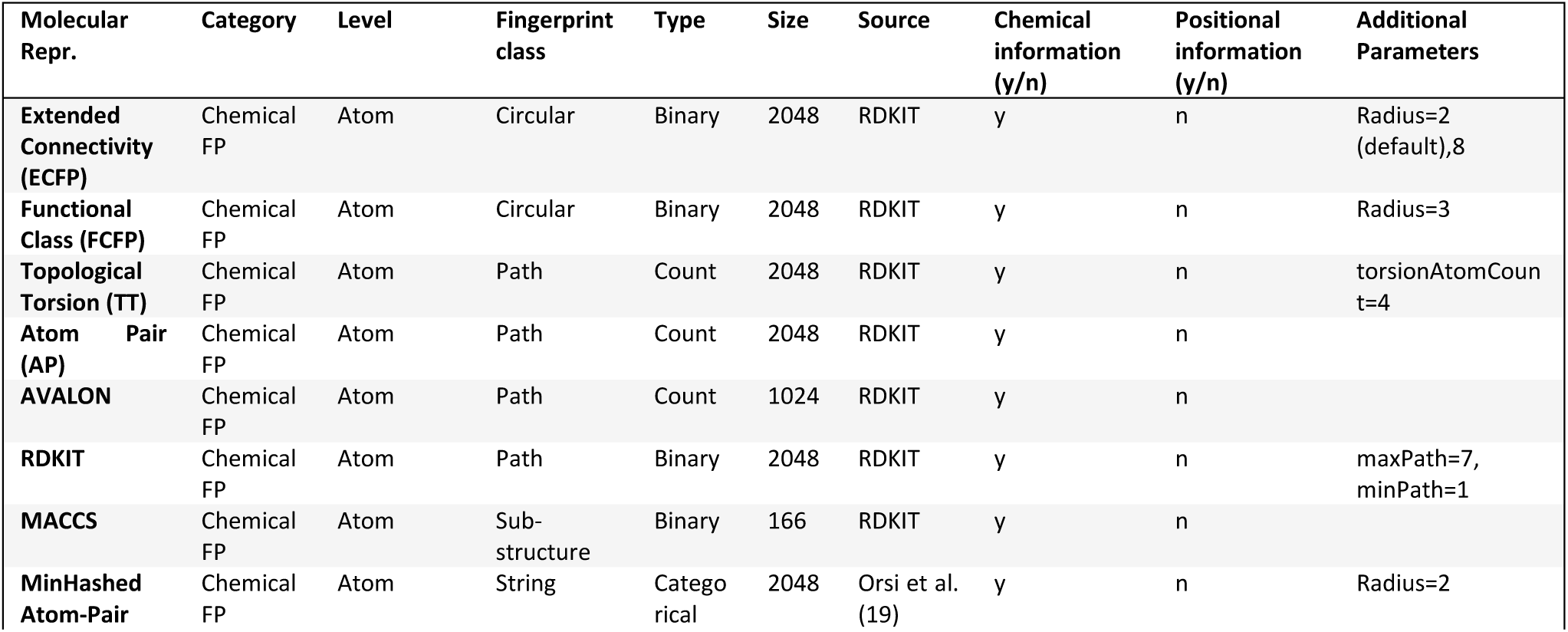
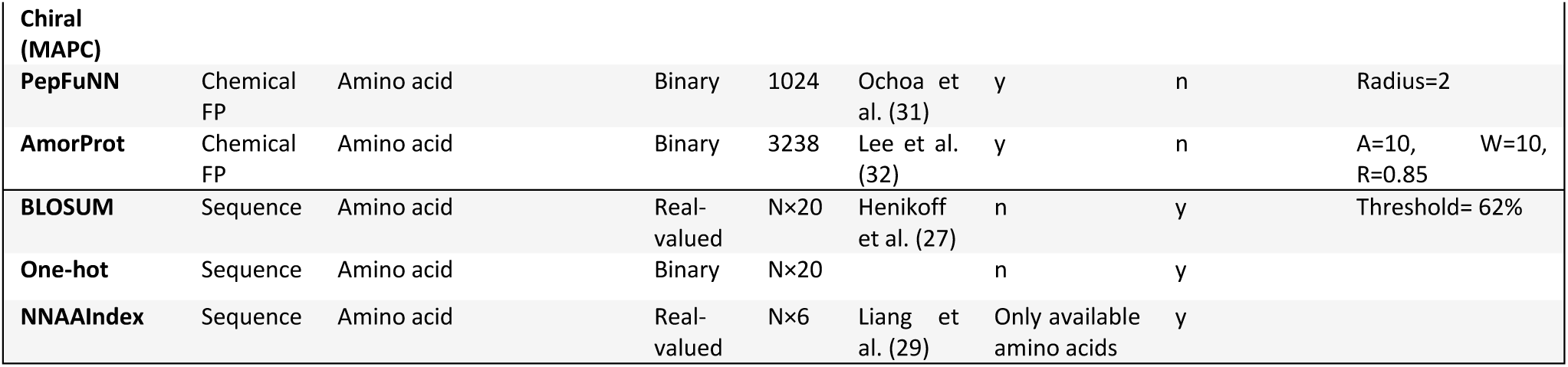
Details of peptide-level chemical fingerprints and sequence-based encodings. Positional information refers to the position of the amino acid residues, and N is the number of amino acids in the peptide sequence.

**Circular fingerprints** describe the local chemical environment around each atom in a molecule up to a specific radius (number of bonds), encoding atom types, bond types and connectivity patterns (18). These features are hashed and aggregated into fixed-length binary or count vectors. We evaluated Extended Connectivity Fingerprints (ECFP) (18) and Functional Class Fingerprints (FCFP) as representative of this class.

**Path-based fingerprints** enumerate the connected atom paths in the molecule and hash them into fixed-length binary or count vectors (30). From this class, we evaluated RDKIT and Avalon.

**Substructure-based fingerprints** encode the presence or absence of predefined structural fragments, including functional groups and specific atom environments. We evaluated MACCS keys, which uses 166 predefined fragments.

**String-based fingerprints** use SMILES string representations as input and apply text-based hashing methods to encode molecular structure (30). We selected MinHashed Atom-Pair Chiral (MAPC) fingerprints, which compute MinHashes from character strings containing SMILES representations of all circular substructure pairs and their topological distances (19).

For all fingerprints, we adopted software-default parameters (**Table 2**). Given the frequent use of Extended Connectivity Fingerprints ECFP16 (Radius = 8) for peptides (21), this variant was also evaluated. We additionally considered fingerprints specifically developed for peptide and protein encoding:

**PepFuNN** (31) is an open-source Novo Nordisk package that builds composition-based fingerprints by converting peptides into graphs with monomers as nodes, calculating tokens from graph-derived fragments, and generating binary fingerprints.

**AmorProt** (32) is a protein sequence representation combining molecular fingerprints of constituent amino acids into a unified peptide fingerprint. Individual amino acid fingerprints (MACCS, ECFP4, ECFP6, and RDKIT) are multiplied by smoothed trigonometric functions, summed column-wise, and normalized to maximum values (32).

#### Direct-encoding fingerprints (residue-level)

The described representations generally capture either chemical or positional information but fail to encompass both properties simultaneously (**Table 2**). To address this limitation, we developed a novel encoding that integrates sequence- and structure-based information, preserving both chemical and positional properties.

Each amino acid, irrespective of whether it is natural or non-natural, can be represented by its SMILES notation and subsequently encoded as a chemical fingerprint. We construct a matrix **F** with dimensions M × N, where each row represents an N-dimensional chemical fingerprint for one of M amino acids in the sequence. By replacing each amino acid with its corresponding fingerprint, we create a position-specific representation that preserves sequence order while incorporating chemical structure information. Since fingerprints describe individual amino acids, we considered a fingerprint length of N=256 for all descriptors, except for MACCS which has a fixed length of N=166 and AmorProt (N = 934).

#### Similarity-based fingerprints (residue-level)

To further explore this concept, we constructed similarity matrices **S** with dimensions M × M, where each entry represents the Tanimoto similarity coefficient between amino acid pairs based on their chemical fingerprints. By encoding pairwise amino acid relationships rather than individual properties, this approach conceptually resembles substitution matrices like BLOSUM and was therefore evaluated alongside direct-encoding fingerprints.

Direct-encoding and similarity-based fingerprints were independently calculated for each considered chemical fingerprint (**Table 2**), yielding 18 distinct matrices for performance comparison. According to the considered fingerprint, we computed Tanimoto similarity as described in Ref. (30). Throughout this work, we denote direct-encoding fingerprints as **F**_fps_ and similarity-based fingerprints as **S**_fps_ where ‘*fps’* indicates the underlying chemical fingerprint (e.g., **F**_ECFP16_, **S**_ECFP16_). We excluded PepFuNN from these representations as it considers the entire peptide for fingerprint calculation. Prior to fingerprint conversion, SMILES strings were canonicalized using RDKit applying charge neutralization and preserving stereochemical information. For consistency across all encoding methods, the BLOSUM62 matrix, one-hot encoding matrix, and similarity matrix **S** are all of dimensions 22 × 22, encompassing the 20 natural amino acids, one post-translationally modified residue (citrullinated arginine), and a placeholder token ‘X’ for unknown residues. For BLOSUM62 and one-hot encoding, we assigned the modified residue and ‘X’ zero scores for all pairwise comparisons. The direct-encoding fingerprints **F** have dimensions 22 x 256 for all fingerprints except for MACCS (22 x 166) and AmorProt (22 x 934). The NNAAIndex matrix has dimensions 22 x 6. We assigned zero scores to ‘X’ for all the obtained matrices.

### 2.3 Neural network model

To test the different descriptors on natural and non-natural amino acids, our machine learning framework builds upon the NNAlign model (14). This method allows predicting the peptide binding affinity and extract the optimal binding core of 9 amino acids. All possible 9-mer windows within each peptide are extracted and processed through a neural network to generate binding scores for each 9-mer. The model then selects the 9-mer with the highest predicted binding score as the predicted binding register for that peptide. This score is compared against the peptide binding affinity score, and the loss is backpropagated to update the network parameters. Through this process, the model learns to identify which 9-mer core within each peptide is most likely responsible for MHC binding. Flanking regions were also considered to capture both the binding register and its context, enabling the neural network to learn binding cores while accounting for the influence of neighbouring residues on pMHCII interactions. For sequence-based encodings, direct-encoding and similarity-based fingerprints, we encoded flanking regions by calculating average scores for sequences surrounding each 9-mer window. The left flank extends backward from the window start, while the right flank extends forward from the window end, with zero-padding applied at sequence boundaries. For peptide-level fingerprints, each flanking region was directly converted to its own fingerprint.

We implemented the neural network model as a feed-forward multilayer perceptron (MLP) with two fully connected hidden layers, each followed by batch normalization (batch size = 32), dropout and rectified linear unit (ReLU) activation function for stabilized and regularized training. Considering that the binding affinity score ranges between 0 and 1, we added sigmoid activation to ensure that the output falls in this range. We employed Adam with weight decay algorithm (33) for stochastic optimization, and, to reduce the potential impact of the large degree of redundancy present in the binding data (14), we modified the network back-propagation so that the step size was divided by the binding core redundancy of the given peptide, as proposed by Nielsen et al. (14). During the core selection phase, the first 15 epochs were considered of ‘build-in’, and only hydrophobic amino acids (‘ILVMFYW’) were enforced to be in position 1. This does not apply for the predictions considering MHC class I alleles and peptide-level fingerprints. We optimized hidden layer dimension, dropout rate, learning rate and weight decay using Optuna (34) (**Table 3**) considering 50 iterations and pruning. For the data sets including NNAA-based peptides, an imbalance factor was further tuned during training to account for imbalance.

**Table 3.**
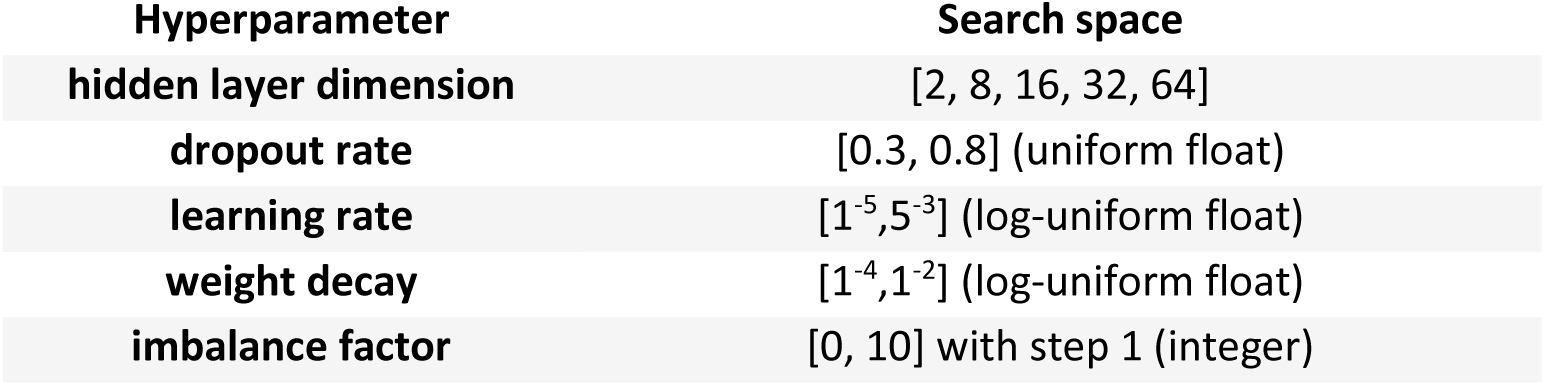
Search space of hyperparameters optimized with Optuna.

As proposed by Nielsen et al. (14), we evaluated a neural network within a nested cross-validation and ensemble framework. We used five-fold outer cross-validation, holding out one fold for evaluation and using the remaining four folds for hyperparameter optimization via a nested four-fold cross-validation. For each Optuna trial, we trained models on three folds and validated on the fourth. Each model was run for 500 epochs, monitoring early stopping (minimum change = 10^−3^, patience = 10) on the validation fold to prevent overfitting. We ranked trials by the mean nested-fold MSE, and the top 5 trials were retained per outer iteration. Each retained trial contributed to its four nested-fold models, yielding an ensemble of up to 20 models per outer fold. We obtained ensemble predictions by averaging model outputs, and binding cores by majority vote across models.

#### Mode of presentation

We adopted two different modes of presentation according to the considered molecular representation. For sequence-based encodings, direct-encoding and similarity-based fingerprints, we encoded each amino acid using the respective descriptor scores, producing concatenated feature vectors of dimension 9 × *d*, where *d* is the descriptor dimensionality. These were concatenated with the average flanking features, yielding final feature vectors with dimension *(9 × d) + 2d*.

For peptide-level fingerprints, the computational cost of calculating fingerprints for each 9-mer window and flanking regions required pre-computation before model training. We processed peptide sequences using a sliding window to extract all possible 9-mer cores, converting each sequence into a single molecular fingerprint. We concatenated core fingerprints with flanking region fingerprints, yielding feature vectors of dimension 3*d*, where *d* is the fingerprint dimensionality (**Table 2**).

### 2.4 Performance evaluation

We used MSE for network training and best model selection. We evaluated model performances through regression lines and coefficient of determination R^2^, root mean square error (RMSE) and Pearson correlation coefficient (PCC). These metrics were computed on the aggregated out-of-fold predictions. In line with studies in immunogenic bioinformatics, we report PCC as main metric for multiple comparisons, reporting R^2^ performances in the Supplementary Information. R^2^ and RMSE are defined as

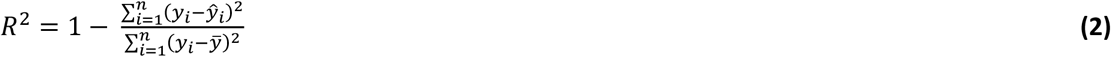

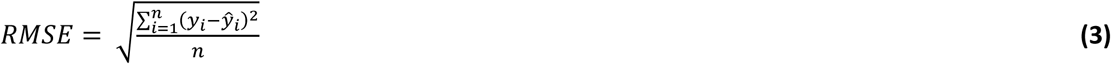

where *n* is the number of samples, *ŷ*_*i*_ is the predicted value of the *i*-th sample, *y*_*i*_ is the corresponding measured value and *ȳ* the mean measured value over *n* samples.

To test for statistically significant differences among molecular representations, we compared each descriptor to BLOSUM62 using paired, per-sample squared error differences computed from out-of-fold predictions using a two-sided Wilcoxon signed-rank test. To account for multiple comparisons across descriptors within each allele, we controlled the false discovery rate at α = 0.05 using the Benjamini-Hochberg procedure, considering significant adjusted p-values < 0.05. We performed all tests on aggregated out-of-fold predictions using the same cross-validation splits across descriptors.

### 2.5 Sequence logos

We calculated sequence logos following the approach presented in Seq2Logo (35), considering Kullback-Leibler divergence, prior weights = 200, and sequence weighting based on a “Hobohm1” like algorithm using an identity threshold of 0.63. For the external test sets derived from the human proteome, sequence logos were constructed from the predicted binding cores of the top 1% strongest predicted binders. For cross-validated predictions, the number of peptides is relatively small (i.e., lower than 6000): all the peptides predicted as binding (ba_score_ ≥ 0.5) were considered for sequence logos, after aggregating estimated binding cores from all out-of-fold predictions.

### 2.6 Evaluating learned binding cores

To evaluate the performance of our method in extracting binding cores, we compared it against two baseline approaches based on TEPITOPE (36), a widely-used benchmark for single-allele MHC class II prediction. The baselines consisted of i) TEPITOPE’s direct predictions, and ii) a two-stage approach where TEPITOPE-extracted cores serve as input to machine learning regressors (MLP, SVM, or XGBoost). This comparison allows evaluating whether cores learned during training by our algorithm can match or exceed the performance achieved using TEPITOPE’s computationally derived cores. We evaluated all approaches on both binding affinity prediction accuracy and the quality of sequence motifs derived from top-1% binders, assessing whether our approach achieves comparable or superior performance to methods relying on pre-defined core extraction algorithms. Additional details regarding this comparison are reported in the Supplementary Information.

### 2.7 Evaluating the importance of encoding chemical information

To test for the importance of encoding chemical information for modified residues, we performed three complementary comparisons. First, to assess model generalization to modified peptides, we trained the models exclusively on natural amino acids and evaluated their predictions on citrullinated peptides. In this comparison, modified residues were encoded as-is for direct-encoding and similarity-based fingerprints, and as ‘X’ tokens or arginine (i.e. citrulline’s closest amino acid analogue) for sequence-based encodings.

Second, to further evaluate generalization abilities and the impact of training data composition for direct-encoding and similarity-based fingerprints, we extended the first approach by systematically varying exposure to citrullinated peptides during training. Starting from the original dataset including natural and modified peptides (**Table 1**), we performed 5-fold cross-validation where each fold served as a test set once. For each cross-validation split, we created five training configurations from the remaining four folds, each containing 0%, 25%, 50%, 75%, or 100% of available citrullinated peptides while retaining all natural peptides, maintaining the ‘Hobohm1’ criterion across both natural and modified peptides to minimize sequence redundancy, as described above. We evaluated performances by aggregating predictions across all five test folds for each training configuration, separately calculating metrics for citrullinated peptides.

Third, we evaluated whether chemical fingerprint encoding improves prediction accuracy when models are trained and tested on datasets that include modified residues. We trained the models using different encoding strategies: i) citrulline, ii) substitution of citrulline with arginine, iii) substitution with ‘X’. Models were trained and evaluated on the same datasets under each encoding strategy. To evaluate the biological relevance of predictions for modified peptides, we analysed the dataset derived from Rebak et al. (26) and assessed the frequency of citrullinated residues across the top 1% strongest predicted binders for each encoding strategy.

## 3. Results

### Direct-encoding and similarity-based fingerprints match sequence encodings for natural peptides

We first evaluated molecular representations based on two criteria: their ability to predict binding affinity scores and their accuracy in assigning binding cores for alleles with experimentally determined cores from 15 HLA-DR restricted peptides compiled from the Protein Data Bank available from Ref. (14).

Peptide-level fingerprints demonstrated critical limitations for pMHCII binding prediction. Overall, these descriptors yielded significantly worse binding affinity predictions than BLOSUM62 (p<0.05), although some descriptors (e.g., AtomPairs fingerprints) performed comparatively better within this class (**Supplementary Figures S1-S2**). More critically, most binding cores (80-100%) were incorrectly identified, even for representations specifically designed for peptides (**Supplementary Figure S3**). By aggregating chemical information across the entire peptide sequence, these descriptors lose the positional information required to identify anchor residues at defined positions. This limitation results in predictions lacking biological interpretability, posing substantial challenges for downstream drug discovery applications.

In contrast, direct-encoding and similarity-based fingerprints correctly identified 80-100% of reference cores (12-14 out of 15 peptides), consistent with sequence-based encoding performance (**Figures 1-2B**). Binding affinity prediction performance varied across descriptors, with NNAAIndex showing the poorest performance overall. The NNAAIndex comprises six representative features encompassing different amino acid properties (29); our results suggest these features are insufficient to achieve performance comparable to BLOSUM62 and other descriptors for pMHCII prediction. Among direct-encoding fingerprints, ECFP4 (**F**_ECFP4_) and AtomPairs (**F**_AP_) demonstrated the best cross-allele performance (**Figure 1A, Supplementary Figure S4**), while MAPC fingerprints (**S**_MAPC_) showed superior overall performance among similarity-based approaches (**Figure 2A, Supplementary Figure S7**). This finding aligns with prior studies, where MAPC descriptors have demonstrated strong performance in several applications, and particularly in similarity tasks (20). The obtained prediction performances generally outperformed TEPITOPE-derived predictions (**Supplementary Figure S12**), consistent with earlier findings (14) that integrated learning of binding cores during training yields competitive or superior performance compared to pre-defined core extraction algorithms.

**Figure 1.**
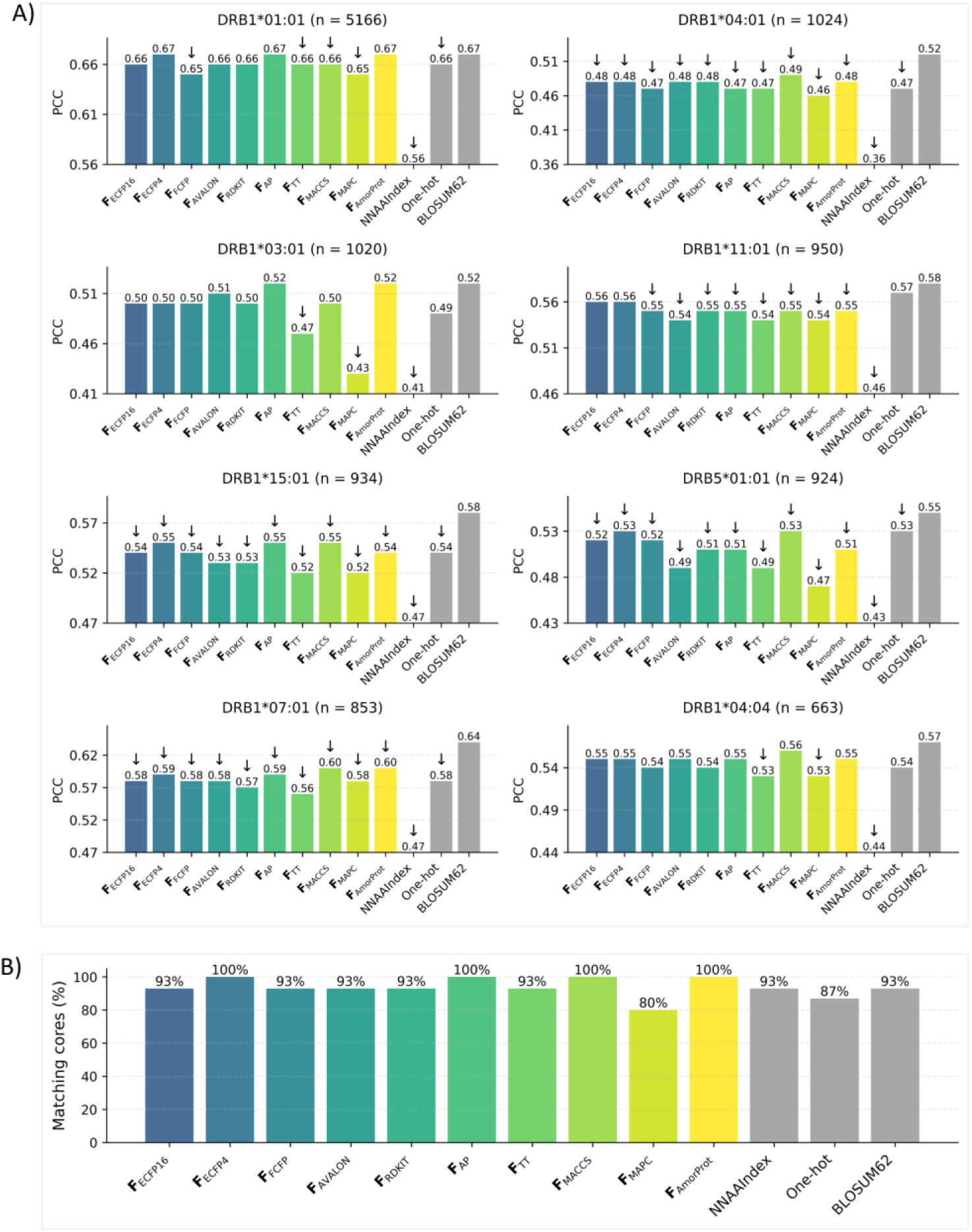
Performance of direct-encoding fingerprints for MHC class II binding prediction. (**A**) PCC values from out-of-fold predictions for natural peptides binding to selected HLA-DR alleles. Each subplot shows results for one allele with sample size indicated in the title. Gray bars represent sequence-based descriptors; coloured bars represent direct-encoding chemical fingerprints. Within each subplot, PCC values are displayed relative to the lowest-performing descriptor to highlight performance differences. Upward arrows indicate significantly better performance than BLOSUM62; downward arrows indicate significantly worse performance than BLOSUM62; absence of arrows indicates no significant difference from BLOSUM62 (paired Wilcoxon tests on per-peptide squared errors, FDR < 0.05, Benjamini-Hochberg correction for 12 pairwise comparisons per allele). Performances for remaining alleles and in terms of R^2^ are reported in **Supplementary Figures S4-S5**. (**B**) Percentage of matching predicted binding cores for each descriptor against reference for 15 HLA-DR restricted peptides compiled from the protein database, retrieved from Ref. (14). 80%, 87% and 93% correspond to 12, 13 and 14 matching cores over 15, respectively. **Supplementary Figure S6** reports detailed information regarding core and peptide sequences.

**Figure 2.**
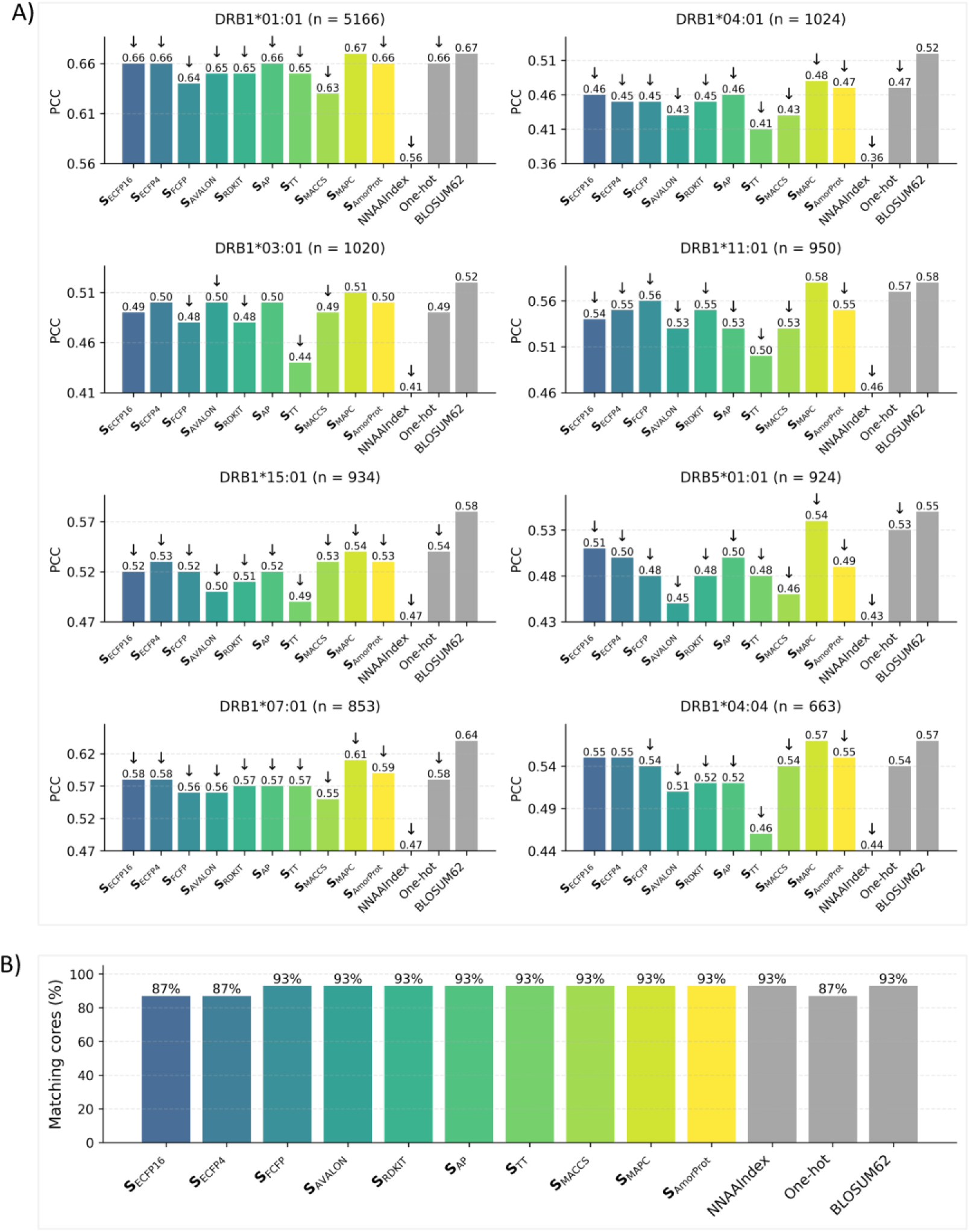
Performance of similarity-based fingerprints for MHC class II binding prediction. (**A**) PCC values from out-of-fold predictions for natural peptides binding to selected HLA-DR alleles. Each subplot shows results for one allele with sample size indicated in the title. Gray bars represent sequence-based descriptors; coloured bars represent similarity-based fingerprints. Within each subplot, PCC values are displayed relative to the lowest-performing descriptor to highlight performance differences. Upward arrows indicate significantly better performance than BLOSUM62; downward arrows indicate significantly worse performance than BLOSUM62; absence of arrows indicates no significant difference from BLOSUM62 (paired Wilcoxon tests on per-peptide squared errors, FDR < 0.05, Benjamini-Hochberg correction for 12 pairwise comparisons per allele). Performances for the remaining alleles and in terms of R^2^ are reported in **Supplementary Figures S7-S8**. (**B**) Percentage of matching predicted binding cores for each descriptor against reference for 15 HLA-DR restricted peptides compiled from the protein database, retrieved from Ref. (14). 87% and 93% correspond to 13 and 14 matching cores over 15, respectively. **Supplementary Figure S9** reports detailed information regarding core and peptide sequences.

MHC class I binding predictions showed similar trends with improved overall performance compared to pMHCII binding prediction (**Figure 3**). Predicting pMHCI binding is typically less challenging than pMHCII, as MHC class I molecules have a closed binding groove that accommodates predominantly 9-mer peptides with specific residues at conserved positions fitting into deep hydrophobic pockets (37). Performance differences across descriptors are less pronounced than for MHC class II predictions, suggesting that the choice of the molecular descriptor becomes less critical for less complex prediction tasks.

**Figure 3.**
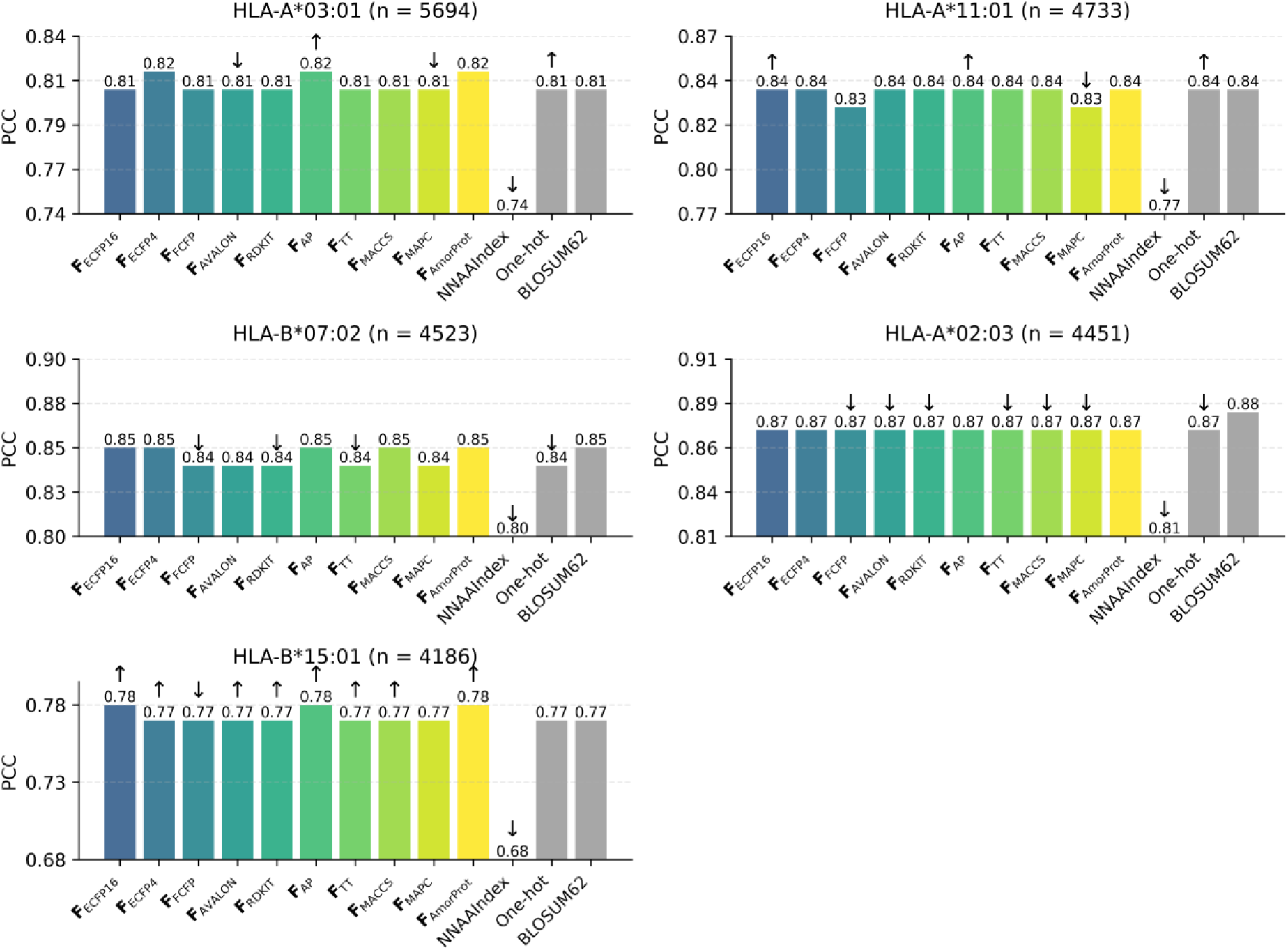
Performance of direct-encoding fingerprints for MHC class I prediction. PCC values from out-of-fold predictions for natural peptides binding to selected HLA-A and HLA-B alleles. Each subplot shows results for one allele with sample size indicated in the title. Gray bars represent sequence-based descriptors; coloured bars represent direct-encoding fingerprints. Within each subplot, PCC values are displayed relative to the lowest-performing descriptor to highlight performance differences. Upward arrows indicate significantly better performance than BLOSUM62; downward arrows indicate significantly worse performance than BLOSUM62; absence of arrows indicates no significant difference from BLOSUM62 (paired Wilcoxon tests on per-peptide squared errors, FDR < 0.05, Benjamini-Hochberg correction for 12 pairwise comparisons per allele). Performances for similarity-based fingerprints and in terms of R^2^ are reported in **Supplementary Figures S14-S16**.

Based on these results, we selected four descriptors representing distinct encoding strategies for subsequent analyses. MAPC similarity-based fingerprints (**S**_MAPC_) demonstrated superior overall performance among similarity-based fingerprints and were selected alongside BLOSUM62 and one-hot encoding, two widely adopted sequence-based encoding methods in immunoinformatics. Among direct-encoding fingerprints, predicted binding scores for the best performing fingerprints showed high correlation (**Supplementary Figure S10**), indicating comparable predictive capability. We selected **F**_AP_ based on its overall better performance across alleles. These four descriptors employ fundamentally different encoding strategies: **F**_AP_ and **S**_MAPC_ capture structural and chemical diversity, one-hot encoding provides categorical amino acid differentiation, and BLOSUM62 incorporates evolutionary substitution information. All four descriptors effectively preserve positional information.

### Predicted binding motifs align with established patterns for natural amino acids

Accurate binding motif prediction is essential for understanding pMHC interactions and immune response mechanisms. **Figure 4** presents predicted binding motifs for representative MHC class II and class I alleles, respectively. For MHC class II, common anchor residues are at positions P1, P4, P6, and P9, with hydrophobic amino acids predominating at P1 (38). The modelled binding motifs align well with the established patterns for HLA-DR allele-binding peptides, characterized by large hydrophobic residues (predominantly phenylalanine and tyrosine) at position 1 and small uncharged residues at position 6 (39). This pattern is evident both in motifs derived from random epitope selection from the human proteome (**Figure 4A**) and out-of-fold predictions (**Supplementary Figure S11**) and remains consistent with TEPITOPE-derived methods (**Supplementary Figure S13**).

**Figure 4.**
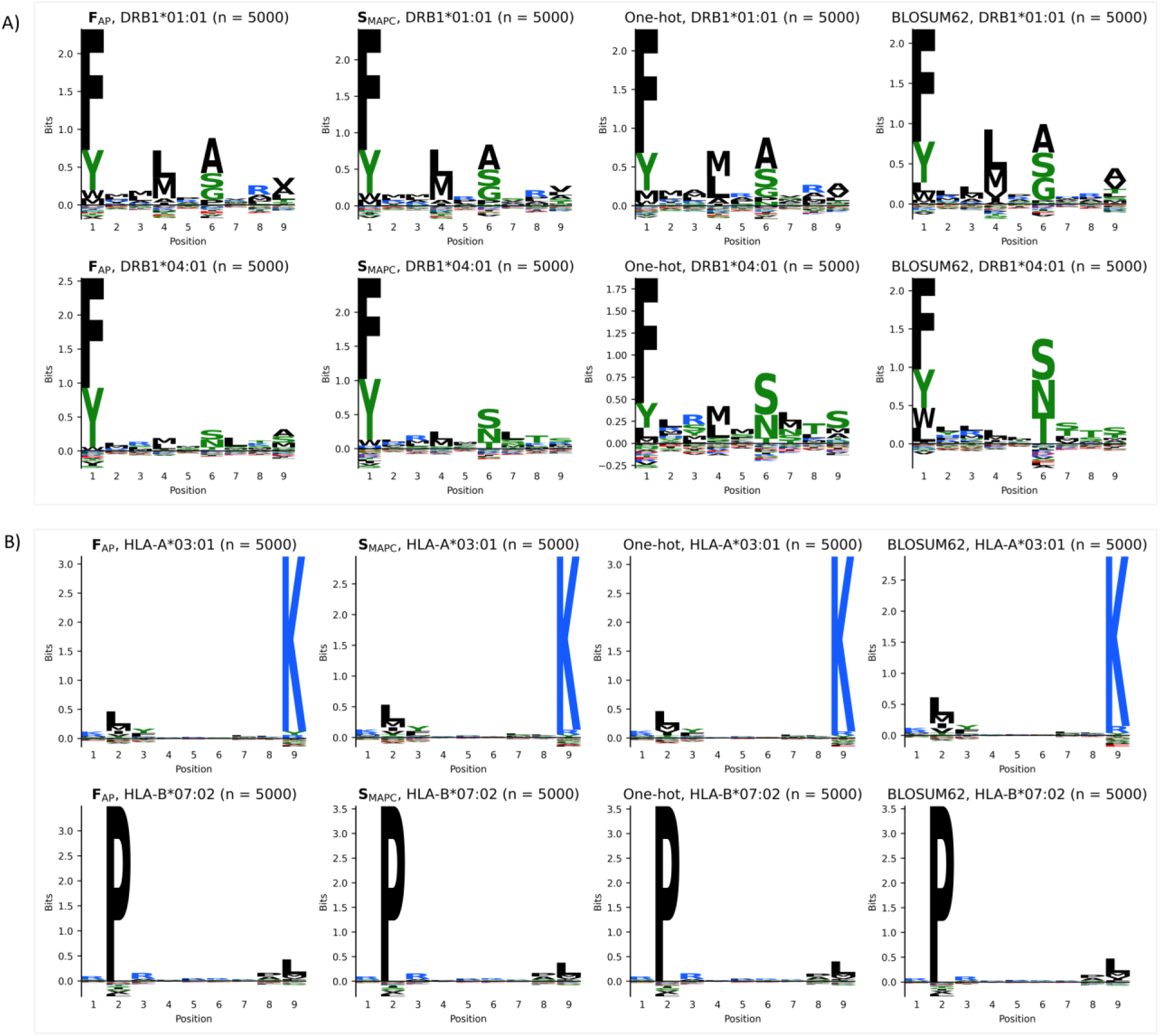
Predicted binding motifs for MHC class II and class I alleles. Sequence logos generated from the top 1% (n = 5000) predicted binders among randomly sampled peptides from the UniProt human proteome for selected (**A**) MHC class II alleles and (**B**) MHC class I alleles, considering **F**_AP_, **S**_MAPC_, one-hot and BLOSUM62 descriptors. Logos display amino acid frequencies at each position, with letter height proportional to information content. Sequence logos from out-of-fold predictions are provided in **Supplementary Figures S11 and S17**.

For MHC class I, primary anchors predominantly occur at positions P2 and P9 of the binding 9-mer (38). **Figure 4B and Supplementary Figure S17** demonstrate that all selected descriptors successfully identify these anchor positions, both for peptides randomly sampled from the human proteome and out-of-fold predictions. HLA-A*03:01 favours leucine and lysine at positions 2 and 9, respectively, consistent with canonical HLA-A3 motifs (40). HLA-B*07:02 shows preference for proline at P2 and leucine at P9, aligning with established patterns (38, 41). These results demonstrate that the proposed descriptors perform comparably to established descriptors for natural amino acids. Testing these descriptors on non-natural amino acids demonstrates their effectiveness for modified peptide binding prediction.

### Chemical encoding improves citrullinated peptide predictions when modifications are included during training

**Figure 5** shows prediction performance for natural and citrullinated peptides binding to DRB1*01:01 and DRB1*04:01 alleles using the four selected encoding strategies. **F**_AP_ and **S**_MAPC_ demonstrate comparable performance across both alleles, though predictions for DRB1*04:01 show reduced accuracy, likely due to limited training data and dataset class imbalance, with fewer binders (ba_score_ ≥ 0.5) compared to non-binders, limiting generalization to binding peptides.

**Figure 5.**
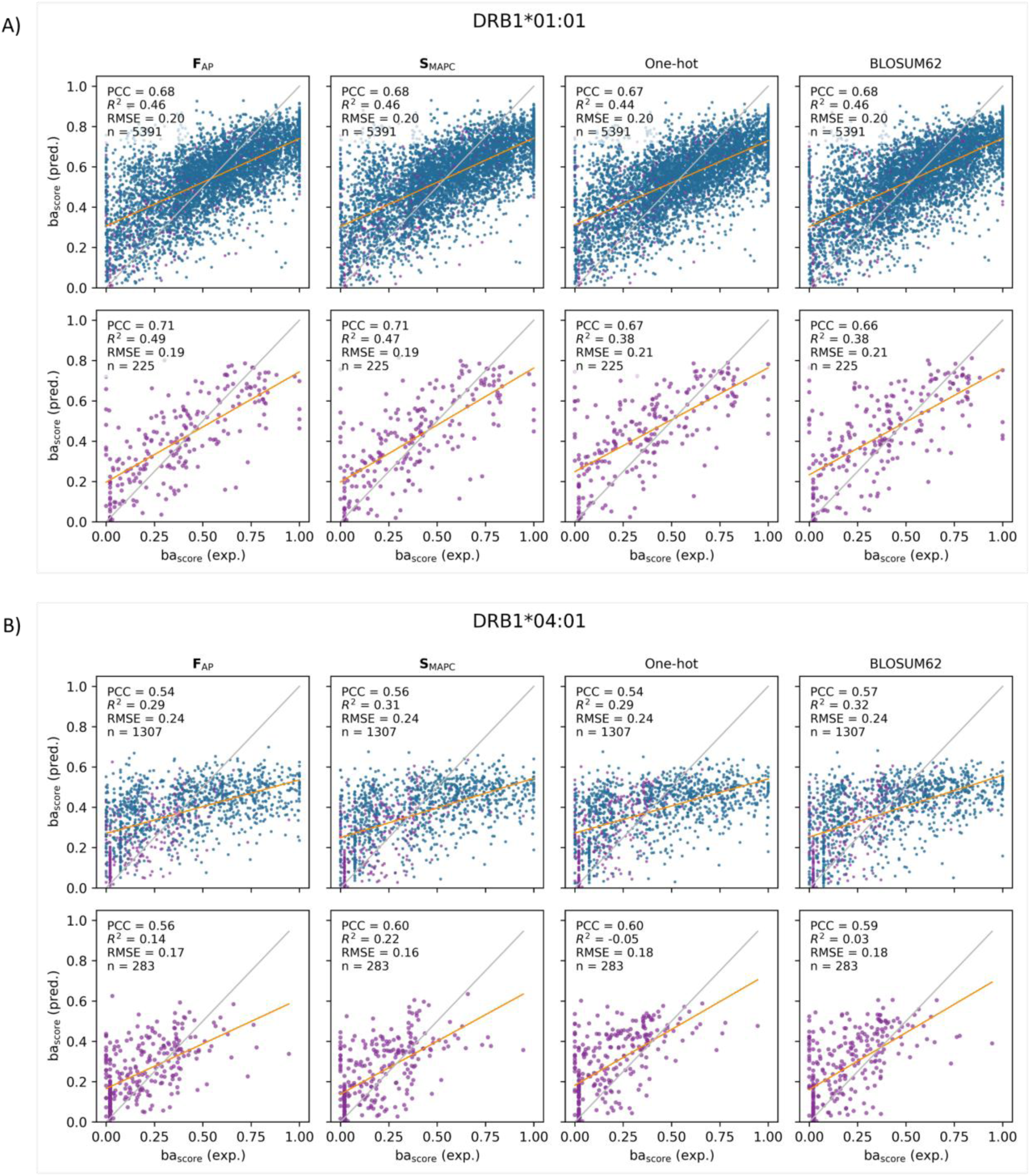
Prediction performance for citrullinated peptides with models trained on natural and modified peptides. Binding affinity predictions for (**A**) DRB1*01:01 and (**B**) DRB1*04:01 using models trained using both natural and citrullinated peptides. In each subplot, the first row shows out-of-fold predictions on both natural and citrullinated peptides, while the second row shows predictions on citrullinated peptides only. Blue dots: natural peptides; purple dots: citrullinated peptides. Yellow line: fitted regression; gray line: identity (perfect prediction). Each panel shows results for four descriptors (**F**_AP_, **S**_MAPC_, one-hot, BLOSUM62) with PCC, R² and RMSE performance metrics. n indicates the number of predicted samples.

While all four descriptors showed comparable performance for natural peptides in both PCC and R² metrics, performance differ for citrullinated peptides, with fingerprint-based encodings (i.e. **F**_AP_ and **S**_MAPC_) showing improved prediction performances in terms of R^2^. This disparity arises because one-hot and BLOSUM62 encode non-canonical residues as zero vectors, disregarding information about modifications during training. PCC values remain relatively comparable across descriptors, indicating that all descriptors capture similar linear relationships between predicted and observed binding affinities. However, R² values differ, with fingerprint-based encodings explaining a greater proportion of variance in binding affinity. This distinction is particularly relevant for applications requiring accurate absolute binding affinity predictions. The preserved linear correlation for one-hot and BLOSUM62 despite zero-vector encoding suggests that models retain a predictive signal through positional information from flanking unmodified residues, though with reduced precision for absolute affinity prediction. As discussed in the following paragraph, evaluating model performance with varying proportions of citrullinated peptides during training and analysing the positional distribution of citrulline in predicted binders provides additional insights into these observations.

### Modifications at non-anchor positions show reduced benefit from explicit chemical encoding

To evaluate the importance of incorporating chemical information for modified amino acids and assess performance generalization to modified peptides, we predicted citrullinated peptides using models trained exclusively on natural peptides (**Figure 6A and Supplementary Figure S18A**). These results show that without encoding citrullination information during training, fingerprint-based descriptors offer no advantage over sequence-based encodings. However, when citrullinated peptides are progressively included during training, R² performance improves for direct-encoding and similarity-based fingerprints starting at 25-50% inclusion, while PCC remains relatively stable across all descriptors (**Figure 6B-C and Supplementary Figures S18B-C**). This indicates that, compared to sequence-based encodings, fingerprint-based encodings can better capture the quantitative contribution of modifications to binding affinity when trained on modified peptides.

**Figure 6.**
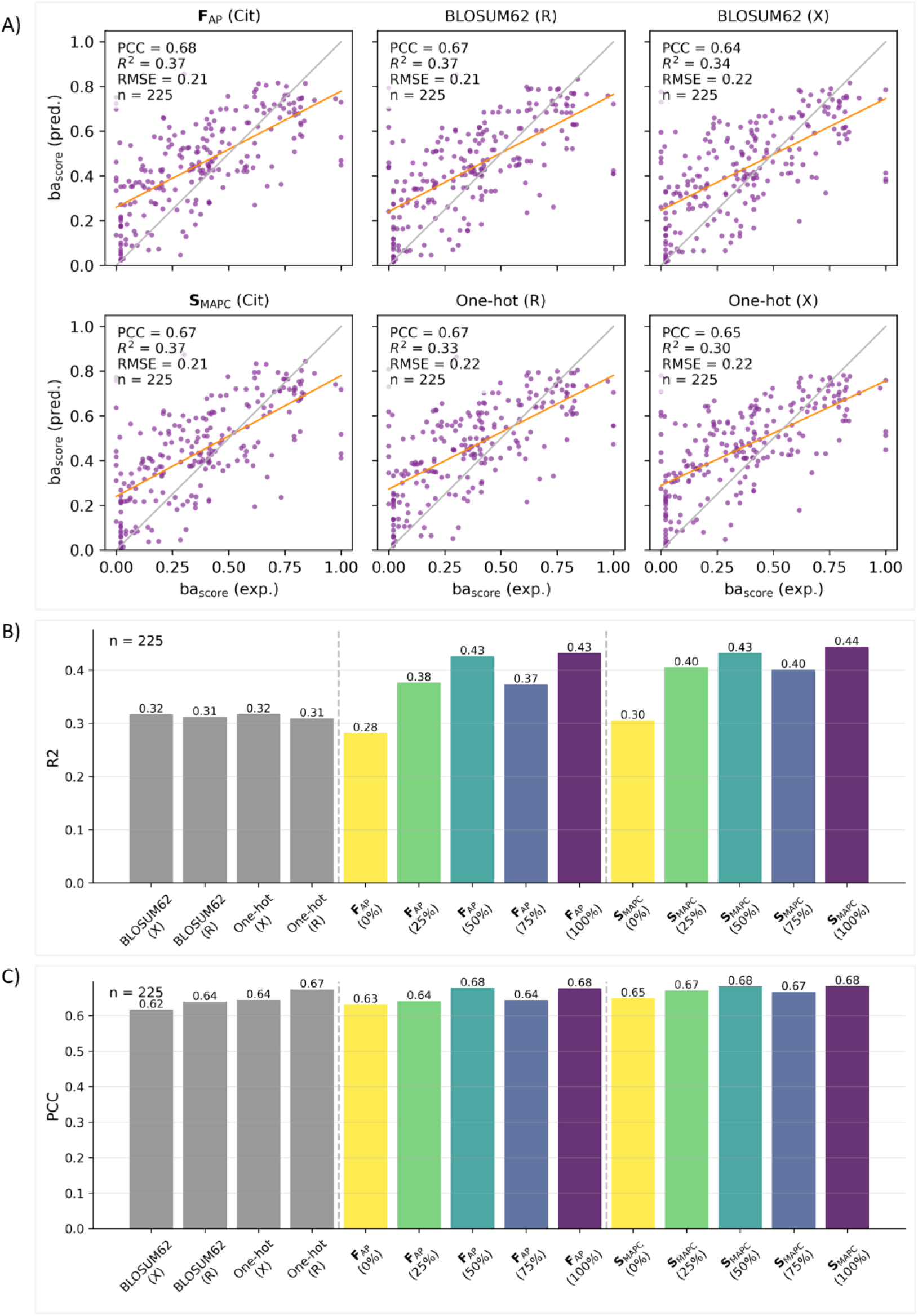
Impact of training data composition on prediction performance for citrullinated peptides (DRB1*01:01). (**A**) Prediction performance on citrullinated peptides using models trained exclusively on canonical peptides. Yellow line: fitted regression; gray line: identity (perfect prediction). Four descriptors are compared: **F**_AP_ and **S**_MAPC_ encode citrulline chemically via fingerprints; one-hot and BLOSUM62 encode citrulline as ‘X’ (zero vector) or ‘R’ (arginine). (**B**) R² and (**C**) PCC as a function of citrullinated peptide proportion in training data (0%, 25%, 50%, 75%, 100%). Models were evaluated using 5-fold cross-validation. Gray bars: BLOSUM62 and one-hot encodings trained only on canonical peptides, with citrulline encoded as ‘X’ or ‘R’; coloured bars: **F**_AP_ and **S**_MAPC_ performance at each training composition. Results for DRB1*04:01 are reported in **Supplementary Figure S18.**

The comparable linear correlation performance (PCC) can be explained by evaluating citrulline frequency in predicted binders (**Figure 7, Supplementary Figure S19**). When citrulline is explicitly encoded with its chemical properties or substituted with arginine, it rarely appears at the four canonical anchor positions (P1, P4, P6, P9). In contrast, when encoded as a zero vector, citrulline shows increased frequency at anchor positions, indicating the model’s inability to distinguish the modification from missing information. These results are in line with earlier findings (42) and suggest that binding affinity is largely determined by unmodified residues at critical anchor positions, allowing sequence-based encodings to achieve linear correlation comparable to fingerprint-based encodings despite lacking explicit citrullination encoding. The need of explicit chemical encoding therefore depends on whether modifications occur at positions critical for binding and whether applications require absolute affinity predictions or only relative comparisons.

**Figure 7.**
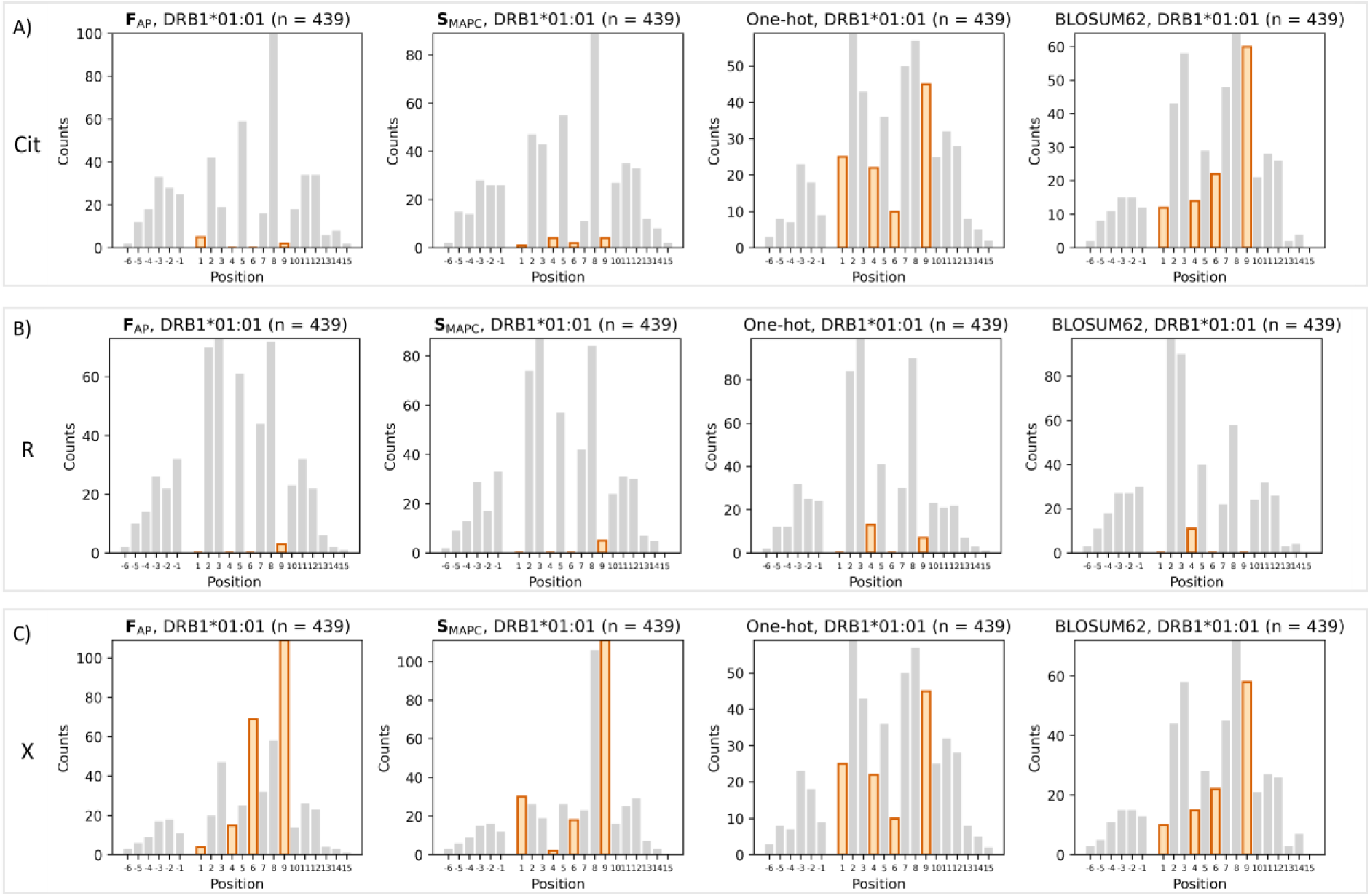
Citrulline positional frequency in predicted binders for DRB1*01:01 under different encoding strategies. Positional occurrence of citrulline in top 1% predicted binders (n = 439) from citrullinated peptides (Rebak et al. (26)), comparing predictions from models trained with citrulline encoded as: (**A**) citrulline (chemical fingerprint for **F**_AP_ and **S**_MAPC_ and zero-vector for one-hot and BLOSUM62), (**B**) arginine (natural analogue), or (**C**) ‘X’ (unknown residue). X-axis shows position relative to the 9-mer binding core: positions 1-9 indicate core positions, negative numbers indicate N-terminal flanking positions, and numbers >9 indicate C-terminal flanking positions. Orange bars highlight frequency at anchor positions (P1, P4, P6, P9); gray bars show frequency at all other positions. Results for DRB1*04:01 are reported in **Supplementary Figure S19**.

### Implementation and availability

The developed models have been included into a user-friendly prediction tool to facilitate immunogenicity assessment on new peptides. Users can predict MHC class II binding affinity for new peptide sequences using the provided prediction script with pre-trained models. The script accepts peptide sequences and supports both canonical and non-canonical amino acids, with non-canonical amino acids encoded in square brackets (e.g., citrulline encoded as [Cit]). After specifying the molecular descriptor and allele, the script outputs predicted binding scores with extracted binding cores for each peptide.

## 4. Discussion

We have presented a machine learning approach combining chemical fingerprints with sequence information to predict MHCII binding affinity for both canonical and modified peptides. A key finding is that residue-level chemical fingerprints (direct-encoding and similarity-based) achieved performances comparable to sequence-based encodings (BLOSUM62 and one-hot) for predicting MHC class I and II binding on natural peptides while accurately identifying binding cores and motifs. In contrast, peptide-level fingerprints demonstrated poor performance due to loss of positional information essential for binding prediction. For citrullinated peptides, chemical encoding substantially improved quantitative prediction accuracy when modified peptides were included during training, demonstrating the importance of explicit chemical representation for more accurate absolute affinity predictions.

Two types of residue-level chemical fingerprints were investigated, showing comparable performance in predicting binding affinity for citrullinated peptides. However, different characteristics should be considered when selecting between these approaches. Direct-encoding fingerprints encode amino acids independently as chemical structures, creating a direct mapping from amino acid positions to their chemical features. In contrast, similarity-based fingerprints encode pairwise relationships between amino acids, requiring an additional computational step to calculate similarity coefficients for all residue pairs. The two representations present substantial differences in dimensionality: for 9-mer cores, direct-encoding fingerprints generate 9 × 256 = 2,304 features, while similarity-based fingerprints produce 9 × 22 = 198 features, scaling with the amino acid vocabulary size. For small datasets, the high feature-to-sample ratio of direct-encoding fingerprints may increase the risk of overfitting, potentially requiring regularization techniques or dimensionality reduction methods. Conversely, direct-encoding fingerprints offer greater flexibility for incorporating non-natural amino acids not encountered during training, which can be directly encoded using their chemical fingerprints. While still possible for similarity-based fingerprints, this requires pre-allocating placeholder positions in the similarity matrix, which are populated with computed similarity scores when new residues are introduced.

Both approaches offer flexibility in accommodating different representations and similarity metrics, enabling future methodological improvements. In this work, we employed the Tanimoto similarity coefficient with diverse chemical fingerprints, yielding a symmetric similarity measure analogous to BLOSUM62. Evaluation of alternative similarity metrics, such as the asymmetric Tversky index, could identify which best captures the chemical relationships relevant to pMHC binding prediction, for instance, the directional nature of amino acid substitutions. Beyond traditional chemical fingerprints, deep learning embeddings have been proposed for pMHC binding prediction (43), and could be integrated as alternative representations to assess their comparative performance for immunogenicity risk assessment.

We tested our descriptors on single-allele pMHC prediction, with a particular focus on MHC class II molecules due to their central role in immunogenicity prediction and therapeutic protein development. However, MHC class II data sets are generally of lower quality compared to MHC class I data, characterized by greater experimental variability and less standardized measurement protocols. Additionally, MHC class II binding predictions are typically more challenging due to the open-ended nature of the binding groove, which accommodates peptides of variable length and permits peptide flanking regions to influence binding. These factors complicate achieving good prediction performances for all alleles. Integrating pan-allele models that incorporate both binding affinity and elution data has demonstrated substantial performance improvements for MHC class II predictions. The employed predictive model and descriptors are compatible with pan-allele frameworks, as they rely on the NNAlign (14) model architecture at the basis of the NetMHCIIpan methods (15). This extension would enable evaluation across a broader range of modifications for which elution data are available in the IEDB.

A critical bottleneck in predicting binding affinity for non-natural amino acids is the scarcity of experimental data. While the IEDB database contains valuable data for biologically relevant post-translational modifications, including phosphorylation and deamidation, we did not incorporate these additional datasets due to several limitations, including lack of or insufficient quantitative binding data, and ambiguous modification annotations. Moreover, these modifications represent only a small fraction of the chemical diversity employed in modern drug design. Medicinal chemists routinely incorporate a vast array of unnatural amino acids into peptide therapeutics to enhance pharmacological properties. Unfortunately, experimental peptide-MHC binding data for these modifications is largely unavailable from public datasets, creating a significant gap in the design of peptide-based therapeutics. Expanding the data set by including a broader range of modifications at various positions, including anchor residues, would improve prediction accuracy and further clarify the impact of chemical encoding on binding affinity. Physics-based computational methods have shown promise in predicting binding affinities for peptide-MHC interactions without relying on experimental training data and will be considered in future work.

By combining concepts of chemical fingerprints and sequence-based encodings, the proposed descriptors represent a paradigm shift in bridging medicinal chemistry and computational immunology, moving beyond sequence-based representations toward explicit chemical property encoding. While current evaluation is limited by available experimental data for modified peptides, the framework’s compatibility with pan-allele models and ability to encode arbitrary chemical modifications enable substantial expansion as data sets become available. This approach addresses a critical gap in immunogenicity prediction tools and can be extended to emerging therapeutic modalities incorporating NNAAs, including therapeutic peptides, antibody-drug conjugates, and synthetic peptide vaccines, enabling immunogenicity risk assessment at the early discovery stage and facilitating mitigation strategies before costly clinical trials.

## 6. Author Contributions

MC: conceptualization, data curation, formal analysis, investigation, methodology, software, validation, visualization, writing-original draft, writing-review and editing. MN: methodology, supervision, writing-review and editing. CB: conceptualization, supervision, writing-review and editing. OO: conceptualization, methodology, funding acquisition, project administration, resources, supervision, writing-review and editing. LDM: conceptualization, methodology, funding acquisition, project administration, resources, supervision, writing-review and editing.

## 7. Acknowledgements

MC is part of the AstraZeneca postdoc program and is grateful for its support. The authors thank Dr. Werngard Czechtizky for the continuous and enthusiastic support given to this project and Dr. Christian Tyrchan, Dr. Eva Nittinger, Dr. Gökce Geylan and Dr. Alessandro Tibo for helpful discussions on the research.

## 8. Generative AI statement

The AI tool Claude Sonnet 4.5 was used for grammar, wording and punctuation review, and during software development. All AI generated text or code was carefully reviewed by the authors before incorporation into the manuscript and Python scripts.

## 9. Data availability

The training dataset including citrullinated peptides will be made publicly available upon manuscript acceptance at the following Zenodo repository: https://doi.org/10.5281/zenodo.19692117.

