## Supplementary Information for "Beyond natural amino acids: Extending immunogenicity risk assessment to non-canonical peptide drugs through chemical feature encoding"

### Contents of this file:

1. Supplementary results for peptide-level fingerprints (Supplementary Figures S1-S3)

2. Supplementary results for direct-encoding and similarity-based fingerprints (pMHCII) (Supplementary Figures S4-S11)

3. Comparison with TEPITOPE-derived methods (Supplementary Figures S12-S13, Supplementary Table S1)

4. Supplementary results for direct-encoding and similarity-based fingerprints (pMHCI) (Supplementary Figures S14-S17)

5. Evaluating the importance of including chemical information (Supplementary Figures S18-S19)

### 1. Supplementary results for peptide-level fingerprints

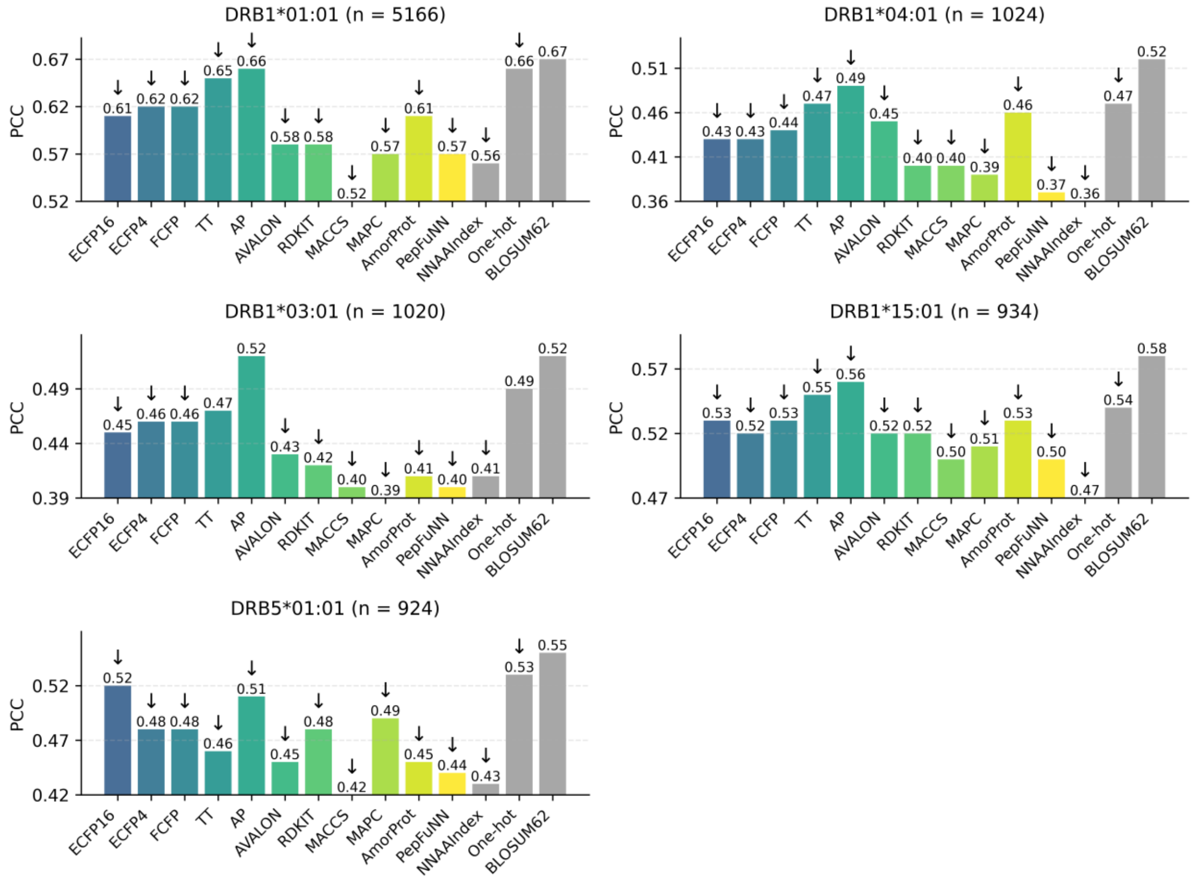


**Supplementary Figure S1. PCC performance of peptide-level fingerprints for MHC class II binding prediction.** PCC values from out-of-fold predictions for natural peptides binding to selected HLA-DR alleles. Each subplot shows results for one allele with sample size indicated in the title. Gray bars represent sequence-based descriptors; coloured bars represent peptide-level chemical fingerprints (see **Table 2** for descriptor details). Within each subplot, PCC values are displayed relative to the lowest-performing descriptor to highlight performance differences. Upward arrows indicate significantly better performance than BLOSUM62; downward arrows indicate significantly worse performance than BLOSUM62; absence of arrows indicates no significant difference from BLOSUM62 (paired Wilcoxon tests on per-peptide squared errors, FDR < 0.05, Benjamini-Hochberg correction for 13 pairwise comparisons per allele).

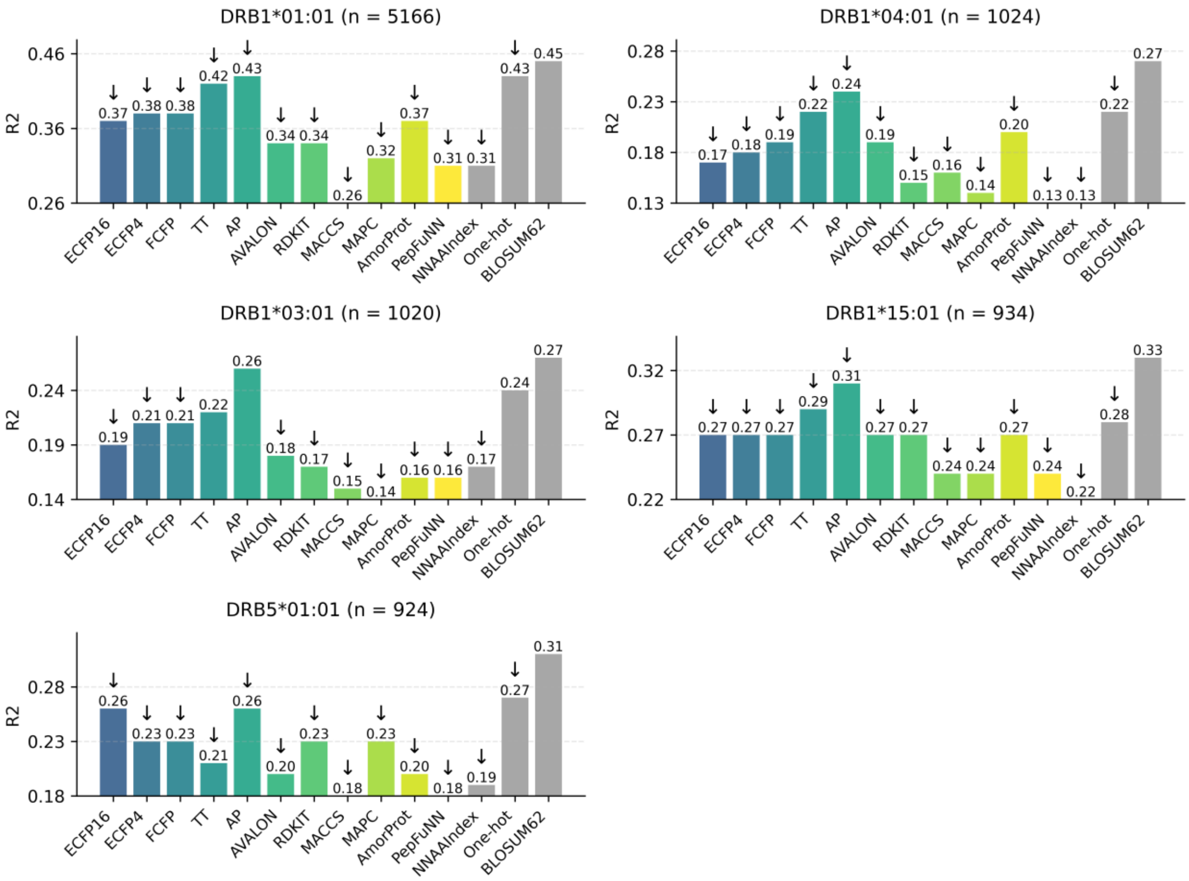


**Supplementary Figure S2. $R^2$ performance of peptide-level fingerprints for MHC class II binding prediction.** $R^2$ values from out-of-fold predictions for natural peptides binding to selected HLA-DR alleles. Each subplot shows results for one allele with sample size indicated in the title. Gray bars represent sequence-based descriptors; coloured bars represent peptide-level chemical fingerprints (see **Table 2** for descriptor details). Within each subplot, $R^2$ values are displayed relative to the lowest-performing descriptor to highlight performance differences. Upward arrows indicate significantly better performance than BLOSUM62; downward arrows indicate significantly worse performance than BLOSUM62; absence of arrows indicates no significant difference from BLOSUM62 (paired Wilcoxon tests on per-peptide squared errors, FDR < 0.05, Benjamini-Hochberg correction for 13 pairwise comparisons per allele).

| Allele | Peptide | Core | ECFP16 | ECFP4 | FCFP | TT | AP | AVALON | RDKIT | MACCS | MAPC | AmorProt | PepFuNN | NNAAIndex | One-hot | BLOSUM62 |
|---|---|---|---|---|---|---|---|---|---|---|---|---|---|---|---|---|
| DRB1*01:01 | AAYSDQATPLLLSPR | YSDQATPLL | QATPLLLSP | QATPLLLSP | DQATPLLLS | QATPLLLSP | DQATPLLLS | QATPLLLSP | DQATPLLLS | QATPLLLSP | SDQATPLLL | QATPLLLSP | QATPLLLSP | YSDQATPLL | YSDQATPLL | YSDQATPLL |
| DRB1*01:01 | AFVKQNAAALA | FVKQNAAAL | VKQNAAALA | VKQNAAALA | VKQNAAALA | VKQNAAALA | VKQNAAALA | VKQNAAALA | VKQNAAALA | VKQNAAALA | AFVKQNAAA | VKQNAAALA | VKQNAAALA | FVKQNAAAL | FVKQNAAAL | FVKQNAAAL |
| DRB1*01:01 | AGFKGEQGPKGEPG | FKGEQGPKG | EQGPKGEPG | GEQGPKGEP | GEQGPKGEP | GEQGPKGEP | GEQGPKGEP | GEQGPKGEP | GEQGPKGEP | GEQGPKGEP | EQGPKGEPG | EQGPKGEPG | GEQGPKGEP | FKGEQGPKG | FKGEQGPKG | FKGEQGPKG |
| DRB1*01:01 | GELIGILNAAKVPAD | IGILNAAKV | ILNAAKVPA | ILNAAKVPA | ILNAAKVPA | ILNAAKVPA | ILNAAKVPA | ILNAAKVPA | ILNAAKVPA | LNAAKVPAD | GELIGILNA | GILNAAKVP | GILNAAKVP | IGILNAAKV | IGILNAAKV | IGILNAAKV |
| DRB1*01:01 | PEVIPMFSALSEGATP | VIPMFSALS | SALSEGATP | SALSEGATP | IPMFSALSE | SALSEGATP | SALSEGATP | IPMFSALSE | SALSEGATP | SALSEGATP | SALSEGATP | SALSEGATP | FSALSEGAT | VIPMFSALS | VIPMFSALS | VIPMFSALS |
| DRB1*01:01 | PKYVKQNTLKLAT | YVKQNTLKL | KQNTLKLAT | KQNTLKLAT | KQNTLKLAT | KQNTLKLAT | VKQNTLKLA | VKQNTLKLA | VKQNTLKLA | KQNTLKLAT | KQNTLKLAT | KQNTLKLAT | KQNTLKLAT | YVKQNTLKL | YVKQNTLKL | YVKQNTLKL |
| DRB1*01:01 | VGSDWRFLRGYHQYA | WRFLRGYHQ | FLRGYHQYA | FLRGYHQYA | FLRGYHQYA | RFLRGYHQY | FLRGYHQYA | RFLRGYHQY | FLRGYHQYA | RFLRGYHQY | FLRGYHQYA | FLRGYHQYA | FLRGYHQYA | WRFLRGYHQ | WRFLRGYHQ | WRFLRGYHQ |
| DRB1*03:01 | PVSKMRMATPLLMQA | MRMATPLLM | VSKMRMATP | VSKMRMATP | VSKMRMATP | PVSKMRMAT | VSKMRMATP | VSKMRMATP | PVSKMRMAT | VSKMRMATP | PVSKMRMAT | MRMATPLLM | RMATPLLMQ | MRMATPLLM | MRMATPLLM | MRMATPLLM |
| DRB1*04:01 | AYMRADAAAGGA | MRADAAAGG | RADAAAGGA | AYMRADAAA | RADAAAGGA | MRADAAAGG | AYMRADAAA | AYMRADAAA | MRADAAAGG | MRADAAAGG | RADAAAGGA | MRADAAAGG | AYMRADAAA | MRADAAAGG | YMRADAAAG | YMRADAAAG |
| DRB1*04:01 | PKYVKQNTLKLAT | YVKQNTLKL | KQNTLKLAT | KQNTLKLAT | KQNTLKLAT | KQNTLKLAT | KQNTLKLAT | KYVKQNTLK | VKQNTLKLA | KQNTLKLAT | KQNTLKLAT | KQNTLKLAT | KQNTLKLAT | YVKQNTLKL | YVKQNTLKL | YVKQNTLKL |
| DRB1*15:01 | ENPVVHFFKNIVTPR | VHFFKNIVT | VHFFKNIVT | VHFFKNIVT | VVHFFKNIV | VHFFKNIVT | VVHFFKNIV | VHFFKNIVT | VHFFKNIVT | VHFFKNIVT | FFKNIVTPR | HFFKNIVTP | VVHFFKNIV | VHFFKNIVT | VHFFKNIVT | VHFFKNIVT |
| DRB1*15:01 | ENPVVHFFKNIVTPRGGSGGGGG | VHFFKNIVT | VVHFFKNIV | VHFFKNIVT | VVHFFKNIV | VHFFKNIVT | VVHFFKNIV | VHFFKNIVT | VHFFKNIVT | VVHFFKNIV | HFFKNIVTP | VVHFFKNIV | VVHFFKNIV | VHFFKNIVT | VHFFKNIVT | VHFFKNIVT |
| DRB5*01:01 | GGVYHFVKKHVHES | YHFVKKHVH | FVKKHVHES | FVKKHVHES | FVKKHVHES | FVKKHVHES | HFVKKHVHE | HFVKKHVHE | HFVKKHVHE | FVKKHVHES | FVKKHVHES | FVKKHVHES | HFVKKHVHE | YHFVKKHVH | FVKKHVHES | YHFVKKHVH |
| DRB5*01:01 | NPVVHFFKNIVTPRTPPPSQ | FKNIVTPRT | IVTPRTPPP | IVTPRTPPP | FKNIVTPRT | KNIVTPRTP | NIVTPRTPP | IVTPRTPPP | IVTPRTPPP | IVTPRTPPP | VHFFKNIVT | IVTPRTPPP | IVTPRTPPP | FKNIVTPRT | FKNIVTPRT | FKNIVTPRT |
| DRB5*01:01 | VHFFKNIVTPRTPGG | FKNIVTPRT | VHFFKNIVT | VHFFKNIVT | NIVTPRTPG | KNIVTPRTP | NIVTPRTPG | IVTPRTPGG | NIVTPRTPG | NIVTPRTPG | IVTPRTPGG | IVTPRTPGG | VHFFKNIVT | FFKNIVTPR | FKNIVTPRT | FKNIVTPRT |
| **Matching** | | | **7%** | **13%** | **7%** | **20%** | **0%** | **13%** | **20%** | **13%** | **0%** | **13%** | **0%** | **93%** | **87%** | **93%** |

**Supplementary Figure S3. Binding core predictions from peptide-level fingerprints compared to experimental references.** Predicted versus reference binding cores for 15 HLA-DR restricted peptides as determined from protein structures (Protein Data Bank, retrieved from Ref. (14)). Columns indicate HLA-DR restriction, peptide sequence, reference core (experimentally determined), and predicted cores for each peptide-level fingerprint. Green: matching predictions; red: divergent predictions.

2. **Supplementary results for direct-encoding and similarity-based fingerprints (pMHCII)**

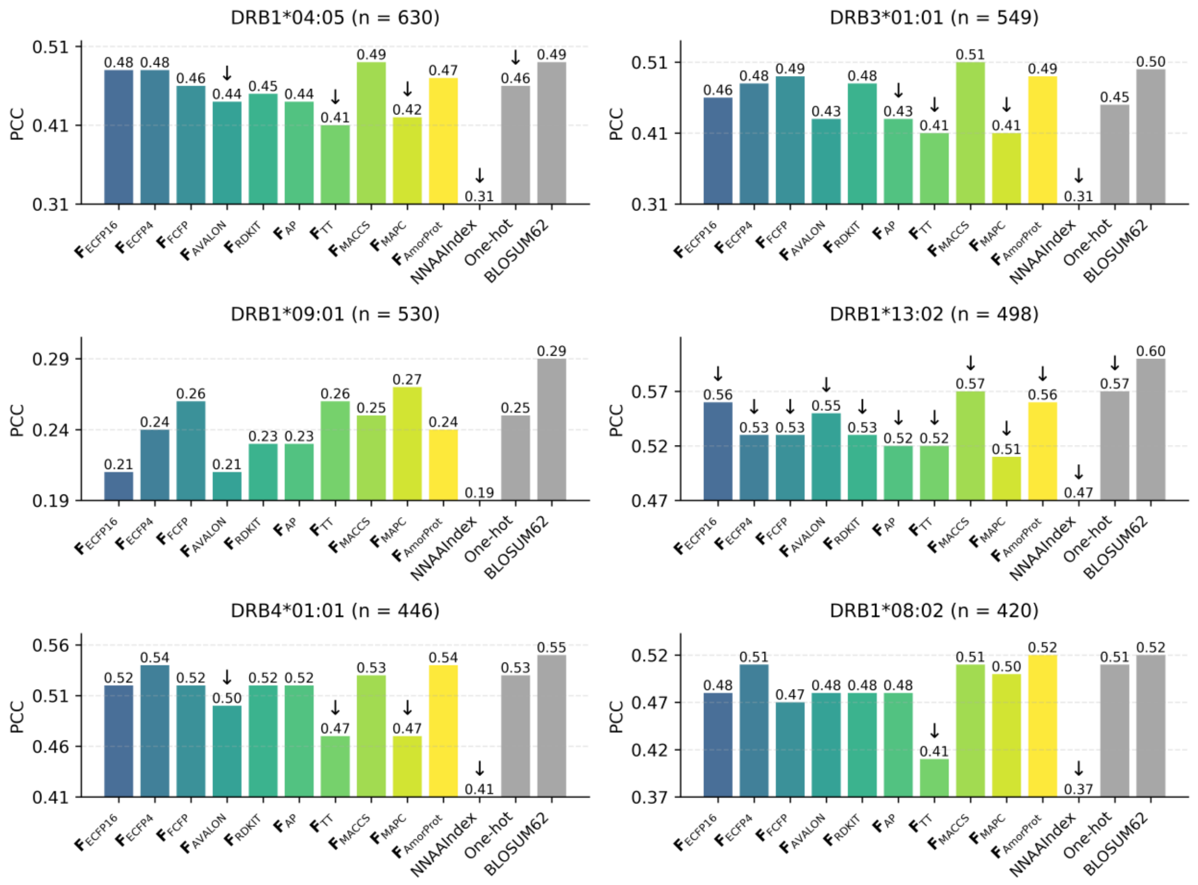


**Supplementary Figure S4. PCC performance of direct-encoding fingerprints for MHC class II binding prediction.** PCC values from out-of-fold predictions for natural peptides binding to selected HLA-DR alleles. Each subplot shows results for one allele with sample size indicated in the title. Gray bars represent sequence-based descriptors; coloured bars represent direct-encoding chemical fingerprints. Within each subplot, PCC values are displayed relative to the lowest-performing descriptor to highlight performance differences. Upward arrows indicate significantly better performance than BLOSUM62; downward arrows indicate significantly worse performance than BLOSUM62; absence of arrows indicates no significant difference from BLOSUM62 (paired Wilcoxon tests on per-peptide squared errors, FDR < 0.05, Benjamini-Hochberg correction for 12 pairwise comparisons per allele).

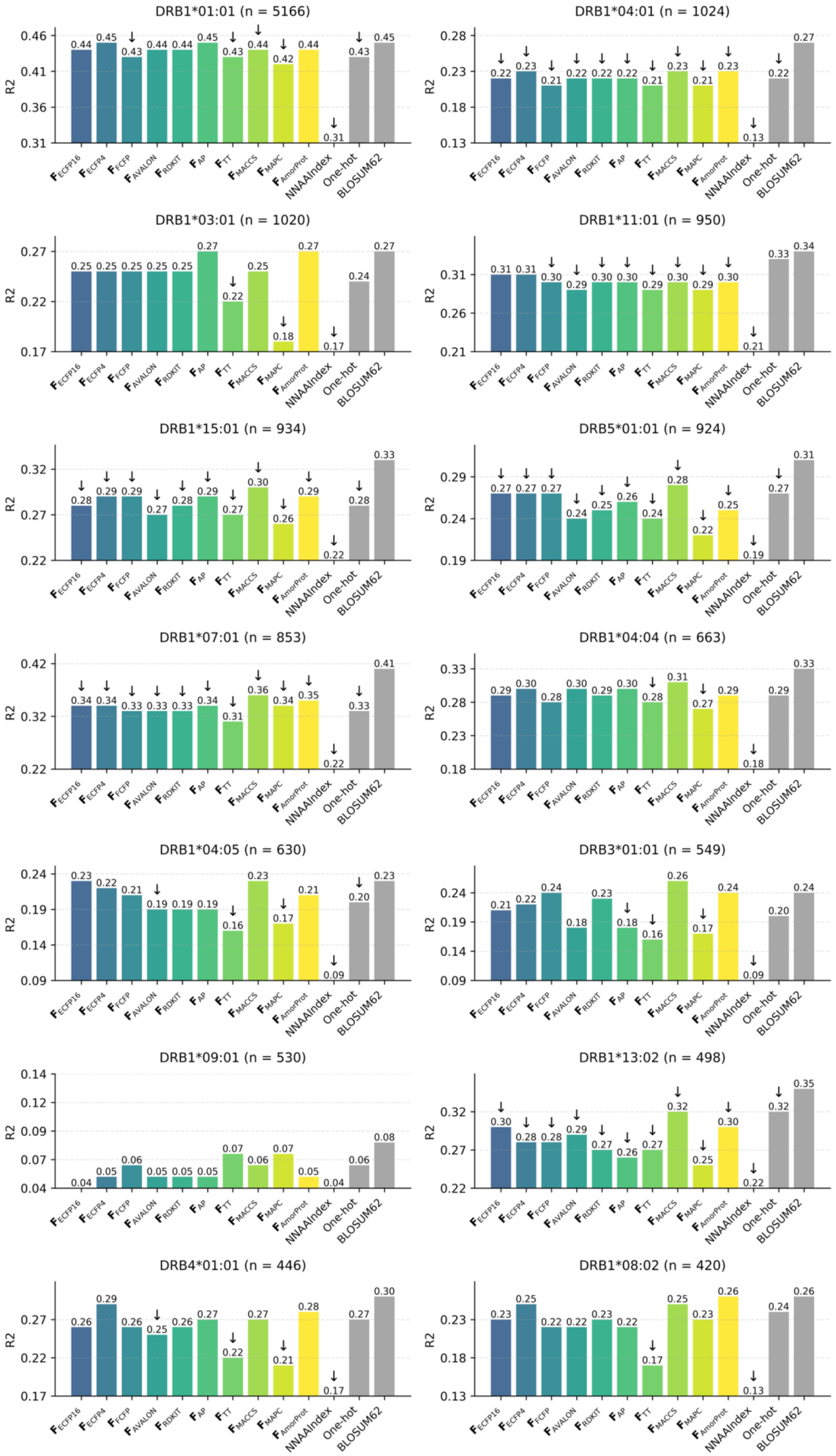

DRB1*01:01 (n = 5166)
R2
0.46 0.41 0.36 0.31
0.44 0.45 0.43 0.44 0.44 0.45 0.43 0.44 0.42 0.44 0.31 0.43 0.45
DRB1*04:01 (n = 1024)
R2
0.28 0.23 0.18 0.13
0.22 0.23 0.21 0.22 0.22 0.22 0.21 0.23 0.21 0.23 0.13 0.22 0.27
DRB1*03:01 (n = 1020)
R2
0.27 0.22 0.17
0.25 0.25 0.25 0.25 0.25 0.27 0.22 0.25 0.18 0.27 0.17 0.24 0.27
DRB1*11:01 (n = 950)
R2
0.31 0.26 0.21
0.31 0.31 0.30 0.29 0.30 0.30 0.29 0.30 0.29 0.30 0.21 0.33 0.34
DRB1*15:01 (n = 934)
R2
0.32 0.27 0.22
0.28 0.29 0.29 0.27 0.28 0.29 0.27 0.30 0.26 0.29 0.22 0.28 0.33
DRB5*01:01 (n = 924)
R2
0.29 0.24 0.19
0.27 0.27 0.27 0.24 0.25 0.26 0.24 0.28 0.22 0.25 0.19 0.27 0.31
DRB1*07:01 (n = 853)
R2
0.42 0.32 0.22
0.34 0.34 0.33 0.33 0.33 0.34 0.31 0.36 0.34 0.35 0.22 0.33 0.41
DRB1*04:04 (n = 663)
R2
0.33 0.28 0.23 0.18
0.29 0.30 0.28 0.30 0.29 0.30 0.28 0.31 0.27 0.29 0.18 0.29 0.33
DRB1*04:05 (n = 630)
R2
0.24 0.19 0.14 0.09
0.23 0.22 0.21 0.19 0.19 0.19 0.16 0.23 0.17 0.21 0.09 0.20 0.23
DRB3*01:01 (n = 549)
R2
0.24 0.19 0.14 0.09
0.21 0.22 0.24 0.18 0.23 0.18 0.16 0.26 0.17 0.24 0.09 0.20 0.24
DRB1*09:01 (n = 530)
R2
0.14 0.12 0.09 0.07 0.04
0.04 0.05 0.06 0.05 0.05 0.05 0.07 0.06 0.07 0.05 0.04 0.06 0.08
DRB1*13:02 (n = 498)
R2
0.32 0.27 0.22
0.30 0.28 0.28 0.29 0.27 0.26 0.27 0.32 0.25 0.30 0.22 0.32 0.35
DRB4*01:01 (n = 446)
R2
0.27 0.22 0.17
0.26 0.29 0.26 0.25 0.26 0.27 0.22 0.27 0.21 0.28 0.17 0.27 0.30
DRB1*08:02 (n = 420)
R2
0.23 0.18 0.13
0.23 0.25 0.22 0.22 0.23 0.22 0.17 0.25 0.23 0.26 0.13 0.24 0.26
$F_{ECFP16}$
$F_{ECFP4}$
$F_{FCFP}$
$F_{AVALON}$
$F_{RDKIT}$
$F_{AP}$
$F_{TT}$
$F_{MACCS}$
$F_{MAPC}$
$F_{AmorProt}$
NNAAIndex
One-hot
BLOSUM62

**Supplementary Figure S5. $R^2$ performance of direct-encoding fingerprints for MHC class II binding prediction.** $R^2$ values from out-of-fold predictions for natural peptides binding to selected HLA-DR alleles. Each subplot shows results for one allele with sample size indicated in the title. Gray bars represent sequence-based descriptors; coloured bars represent direct-encoding chemical fingerprints. Within each subplot, $R^2$ values are displayed relative to the lowest-performing descriptor to highlight performance differences. Upward arrows indicate significantly better performance than BLOSUM62; downward arrows indicate significantly worse performance than BLOSUM62; absence of arrows indicates no significant difference from BLOSUM62 (paired Wilcoxon tests on per-peptide squared errors, FDR < 0.05, Benjamini-Hochberg correction for 12 pairwise comparisons per allele).

| Allele | Peptide | Core | $F_{ECFP16}$ | $F_{ECFP4}$ | $F_{FCFP}$ | $F_{AVALON}$ | $F_{RDKIT}$ | $F_{AP}$ | $F_{TT}$ | $F_{MACCS}$ | $F_{MAPC}$ | $F_{AmorProt}$ | NNAAIndex | One-hot | BLOSUM62 |
|---|---|---|---|---|---|---|---|---|---|---|---|---|---|---|---|
| DRB1*01:01 | AAYSDQATPLLLSPR | YSDQATPLL | YSDQATPLL | YSDQATPLL | YSDQATPLL | YSDQATPLL | YSDQATPLL | YSDQATPLL | YSDQATPLL | YSDQATPLL | YSDQATPLL | YSDQATPLL | YSDQATPLL | YSDQATPLL | YSDQATPLL |
| DRB1*01:01 | AFVKQNAAALA | FVKQNAAAL | FVKQNAAAL | FVKQNAAAL | FVKQNAAAL | FVKQNAAAL | FVKQNAAAL | FVKQNAAAL | FVKQNAAAL | FVKQNAAAL | FVKQNAAAL | FVKQNAAAL | FVKQNAAAL | FVKQNAAAL | FVKQNAAAL |
| DRB1*01:01 | AGFKGEQGPKGEPG | FKGEQGPKG | FKGEQGPKG | FKGEQGPKG | FKGEQGPKG | FKGEQGPKG | FKGEQGPKG | FKGEQGPKG | FKGEQGPKG | FKGEQGPKG | FKGEQGPKG | FKGEQGPKG | FKGEQGPKG | FKGEQGPKG | FKGEQGPKG |
| DRB1*01:01 | GELIGILNAAKVPAD | IGILNAAKV | IGILNAAKV | IGILNAAKV | IGILNAAKV | IGILNAAKV | IGILNAAKV | IGILNAAKV | IGILNAAKV | IGILNAAKV | IGILNAAKV | IGILNAAKV | IGILNAAKV | IGILNAAKV | IGILNAAKV |
| DRB1*01:01 | PEVIPMFSALSEGATP | VIPMFSALS | VIPMFSALS | VIPMFSALS | VIPMFSALS | VIPMFSALS | VIPMFSALS | VIPMFSALS | VIPMFSALS | VIPMFSALS | VIPMFSALS | VIPMFSALS | VIPMFSALS | VIPMFSALS | VIPMFSALS |
| DRB1*01:01 | PKYVKQNTLKLAT | YVKQNTLKL | YVKQNTLKL | YVKQNTLKL | YVKQNTLKL | YVKQNTLKL | YVKQNTLKL | YVKQNTLKL | YVKQNTLKL | YVKQNTLKL | YVKQNTLKL | YVKQNTLKL | YVKQNTLKL | YVKQNTLKL | YVKQNTLKL |
| DRB1*01:01 | VGSDWRFLRGYHQYA | WRFLRGYHQ | WRFLRGYHQ | WRFLRGYHQ | WRFLRGYHQ | WRFLRGYHQ | WRFLRGYHQ | WRFLRGYHQ | WRFLRGYHQ | WRFLRGYHQ | WRFLRGYHQ | WRFLRGYHQ | WRFLRGYHQ | WRFLRGYHQ | WRFLRGYHQ |
| DRB1*03:01 | PVSKMRMATPLLMQA | MRMATPLLM | MRMATPLLM | MRMATPLLM | MRMATPLLM | MRMATPLLM | MRMATPLLM | MRMATPLLM | MRMATPLLM | MRMATPLLM | MRMATPLLM | MRMATPLLM | MRMATPLLM | MRMATPLLM | MRMATPLLM |
| DRB1*04:01 | AYMRADAAAGGA | MRADAAAGG | MRADAAAGG | MRADAAAGG | YMRADAAAG | YMRADAAAG | YMRADAAAG | MRADAAAGG | MRADAAAGG | MRADAAAGG | YMRADAAAG | MRADAAAGG | MRADAAAGG | YMRADAAAG | YMRADAAAG |
| DRB1*04:01 | PKYVKQNTLKLAT | YVKQNTLKL | YVKQNTLKL | YVKQNTLKL | YVKQNTLKL | YVKQNTLKL | YVKQNTLKL | YVKQNTLKL | YVKQNTLKL | YVKQNTLKL | YVKQNTLKL | YVKQNTLKL | YVKQNTLKL | YVKQNTLKL | YVKQNTLKL |
| DRB1*15:01 | ENPVVHFFKNIVTPR | VHFFKNIVT | VHFFKNIVT | VHFFKNIVT | VHFFKNIVT | VHFFKNIVT | VHFFKNIVT | VHFFKNIVT | VHFFKNIVT | VHFFKNIVT | VHFFKNIVT | VHFFKNIVT | VHFFKNIVT | VHFFKNIVT | VHFFKNIVT |
| DRB1*15:01 | ENPVVHFFKNIVTPRGGSGGGGG | VHFFKNIVT | VHFFKNIVT | VHFFKNIVT | VHFFKNIVT | VHFFKNIVT | VHFFKNIVT | VHFFKNIVT | VHFFKNIVT | VHFFKNIVT | VHFFKNIVT | VHFFKNIVT | VHFFKNIVT | VHFFKNIVT | VHFFKNIVT |
| DRB5*01:01 | GGVYHFVKKHVHES | YHFVKKHVH | FVKKHVHES | YHFVKKHVH | YHFVKKHVH | YHFVKKHVH | YHFVKKHVH | YHFVKKHVH | FVKKHVHES | YHFVKKHVH | YHFVKKHVH | YHFVKKHVH | YHFVKKHVH | FVKKHVHES | YHFVKKHVH |
| DRB5*01:01 | NPVVHFFKNIVTPRTPPPSQ | FKNIVTPRT | FKNIVTPRT | FKNIVTPRT | FKNIVTPRT | FKNIVTPRT | FKNIVTPRT | FKNIVTPRT | FKNIVTPRT | FKNIVTPRT | FFKNIVTPR | FKNIVTPRT | FKNIVTPRT | FKNIVTPRT | FKNIVTPRT |
| DRB5*01:01 | VHFFKNIVTPRTPGG | FKNIVTPRT | FKNIVTPRT | FKNIVTPRT | FKNIVTPRT | FKNIVTPRT | FKNIVTPRT | FKNIVTPRT | FKNIVTPRT | FKNIVTPRT | FFKNIVTPR | FKNIVTPRT | FFKNIVTPR | FKNIVTPRT | FKNIVTPRT |
| Matching | | | 93% | 100% | 93% | 93% | 93% | 100% | 93% | 100% | 80% | 100% | 93% | 87% | 93% |

**Supplementary Figure S6. Binding core predictions from direct-encoding fingerprints compared to experimental references.** Predicted versus reference binding cores for 15 HLA-DR restricted peptides as determined from protein structures (Protein Data Bank, retrieved from Ref. (14)). Columns indicate HLA-DR restriction, peptide sequence, reference core (experimentally determined), and predicted cores for each peptide-level fingerprint. Green: matching predictions; red: divergent predictions.

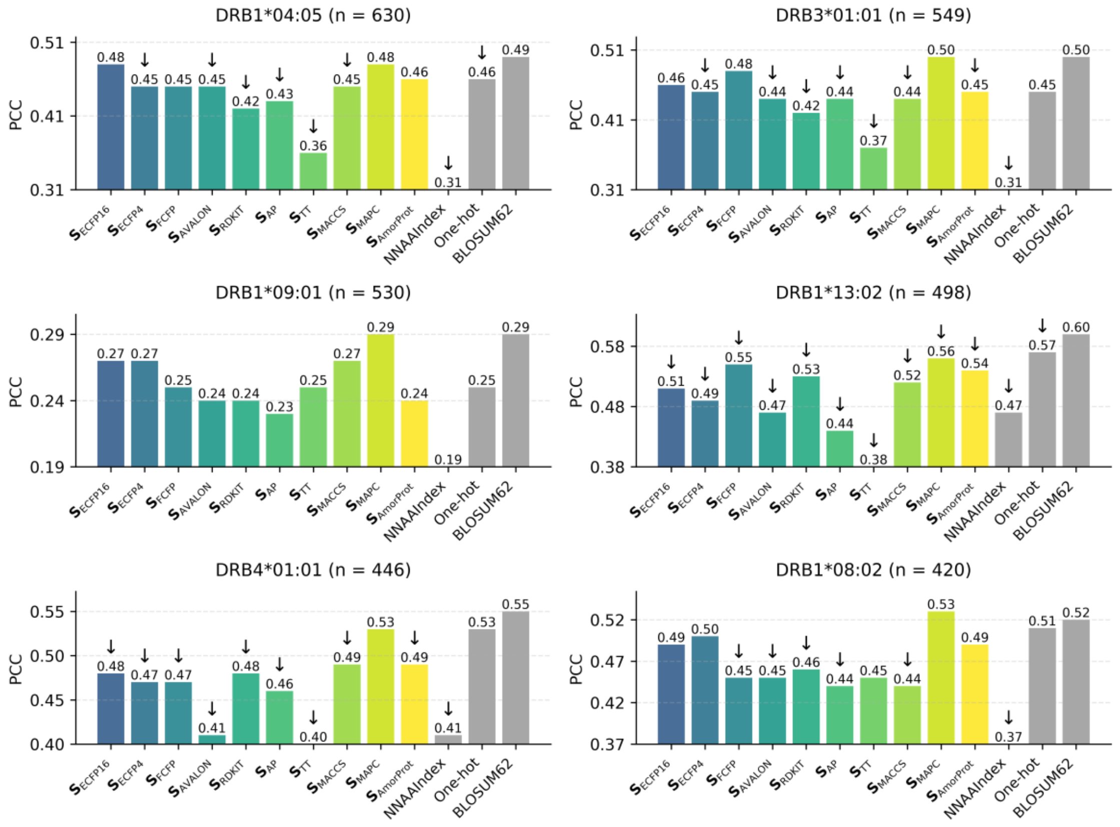


**Supplementary Figure S7. PCC performance of similarity-based fingerprints for MHC class II binding prediction.** PCC values from out-of-fold predictions for natural peptides binding to selected HLA-DR alleles. Each subplot shows results for one allele with sample size indicated in the title. Gray bars represent sequence-based descriptors; coloured bars represent similarity-based fingerprints. Within each subplot, PCC values are displayed relative to the lowest-performing descriptor to highlight performance differences. Upward arrows indicate significantly better performance than BLOSUM62; downward arrows indicate significantly worse performance than BLOSUM62; absence of arrows indicates no significant difference from BLOSUM62 (paired Wilcoxon tests on per-peptide squared errors, FDR < 0.05, Benjamini-Hochberg correction for 12 pairwise comparisons per allele).

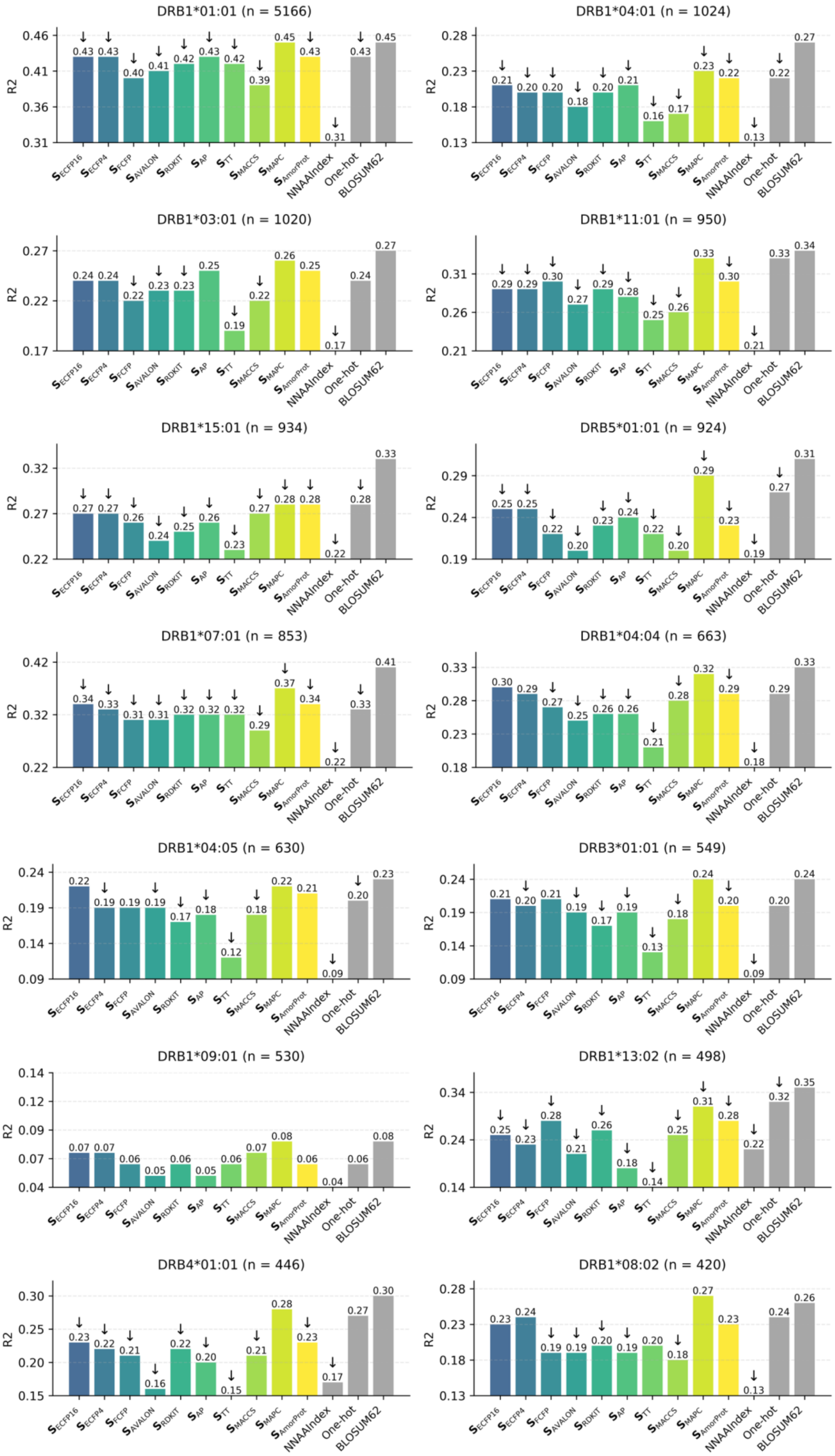
DRB1*01:01 (n = 5166)
DRB1*04:01 (n = 1024)
DRB1*03:01 (n = 1020)
DRB1*11:01 (n = 950)
DRB1*15:01 (n = 934)
DRB5*01:01 (n = 924)
DRB1*07:01 (n = 853)
DRB1*04:04 (n = 663)
DRB1*04:05 (n = 630)
DRB3*01:01 (n = 549)
DRB1*09:01 (n = 530)
DRB1*13:02 (n = 498)
DRB4*01:01 (n = 446)
DRB1*08:02 (n = 420)
R2
S ECFP16
S ECFP4
S FCFP
S AVALON
S RDKIT
S AP
S TT
S MACCS
S MAPC
S AmorProt
NNAAIndex
One-hot
BLOSUM62

**Supplementary Figure S8. $R^2$ performance of similarity-based fingerprints for MHC class II binding prediction.** $R^2$ values from out-of-fold predictions for natural peptides binding to selected HLA-DR alleles. Each subplot shows results for one allele with sample size indicated in the title. Gray bars represent sequence-based descriptors; coloured bars represent similarity-based fingerprints. Within each subplot, $R^2$ values are displayed relative to the lowest-performing descriptor to highlight performance differences. Upward arrows indicate significantly better performance than BLOSUM62; downward arrows indicate significantly worse performance than BLOSUM62; absence of arrows indicates no significant difference from BLOSUM62 (paired Wilcoxon tests on per-peptide squared errors, FDR < 0.05, Benjamini-Hochberg correction for 12 pairwise comparisons per allele).

| Allele | Peptide | Core | $S_{ECFP16}$ | $S_{ECFP4}$ | $S_{FCFP}$ | $S_{AVALON}$ | $S_{RDKIT}$ | $S_{AP}$ | $S_{TT}$ | $S_{MACCS}$ | $S_{MAPC}$ | $S_{AmorProt}$ | NNAAIndex | One-hot | BLOSUM62 |
|---|---|---|---|---|---|---|---|---|---|---|---|---|---|---|---|
| DRB1*01:01 | AAYSDQATPLLLSPR | YSDQATPLL | YSDQATPLL | YSDQATPLL | YSDQATPLL | YSDQATPLL | YSDQATPLL | YSDQATPLL | YSDQATPLL | YSDQATPLL | YSDQATPLL | YSDQATPLL | YSDQATPLL | YSDQATPLL | YSDQATPLL |
| DRB1*01:01 | AFVKQNAAALA | FVKQNAAAL | FVKQNAAAL | FVKQNAAAL | FVKQNAAAL | FVKQNAAAL | FVKQNAAAL | FVKQNAAAL | FVKQNAAAL | FVKQNAAAL | FVKQNAAAL | FVKQNAAAL | FVKQNAAAL | FVKQNAAAL | FVKQNAAAL |
| DRB1*01:01 | AGFKGEQGPKGEPG | FKGEQGPKG | FKGEQGPKG | FKGEQGPKG | FKGEQGPKG | FKGEQGPKG | FKGEQGPKG | FKGEQGPKG | FKGEQGPKG | FKGEQGPKG | FKGEQGPKG | FKGEQGPKG | FKGEQGPKG | FKGEQGPKG | FKGEQGPKG |
| DRB1*01:01 | GELIGILNAAKVPAD | IGILNAAKV | IGILNAAKV | IGILNAAKV | IGILNAAKV | IGILNAAKV | IGILNAAKV | IGILNAAKV | IGILNAAKV | IGILNAAKV | IGILNAAKV | IGILNAAKV | IGILNAAKV | IGILNAAKV | IGILNAAKV |
| DRB1*01:01 | PEVIPMFSALSEGATP | VIPMFSALS | VIPMFSALS | VIPMFSALS | VIPMFSALS | VIPMFSALS | VIPMFSALS | VIPMFSALS | VIPMFSALS | VIPMFSALS | VIPMFSALS | VIPMFSALS | VIPMFSALS | VIPMFSALS | VIPMFSALS |
| DRB1*01:01 | PKYVKQNTLKLAT | YVKQNTLKL | YVKQNTLKL | YVKQNTLKL | YVKQNTLKL | YVKQNTLKL | YVKQNTLKL | YVKQNTLKL | YVKQNTLKL | YVKQNTLKL | YVKQNTLKL | YVKQNTLKL | YVKQNTLKL | YVKQNTLKL | YVKQNTLKL |
| DRB1*01:01 | VGSDWRFLRGYHQYA | WRFLRGYHQ | WRFLRGYHQ | WRFLRGYHQ | WRFLRGYHQ | WRFLRGYHQ | WRFLRGYHQ | WRFLRGYHQ | WRFLRGYHQ | WRFLRGYHQ | WRFLRGYHQ | WRFLRGYHQ | WRFLRGYHQ | WRFLRGYHQ | WRFLRGYHQ |
| DRB1*03:01 | PVSKMRMATPLLMQA | MRMATPLLM | MRMATPLLM | MRMATPLLM | MRMATPLLM | MRMATPLLM | MRMATPLLM | MRMATPLLM | MRMATPLLM | MRMATPLLM | MRMATPLLM | MRMATPLLM | MRMATPLLM | MRMATPLLM | MRMATPLLM |
| DRB1*04:01 | AYMRADAAAGGA | MRADAAAGG | YMRADAAAG | YMRADAAAG | YMRADAAAG | YMRADAAAG | YMRADAAAG | YMRADAAAG | YMRADAAAG | YMRADAAAG | YMRADAAAG | YMRADAAAG | MRADAAAGG | YMRADAAAG | YMRADAAAG |
| DRB1*04:01 | PKYVKQNTLKLAT | YVKQNTLKL | YVKQNTLKL | YVKQNTLKL | YVKQNTLKL | YVKQNTLKL | YVKQNTLKL | YVKQNTLKL | YVKQNTLKL | YVKQNTLKL | YVKQNTLKL | YVKQNTLKL | YVKQNTLKL | YVKQNTLKL | YVKQNTLKL |
| DRB1*15:01 | ENPVVHFFKNIVTPR | VHFFKNIVT | FFKNIVTPR | FFKNIVTPR | VHFFKNIVT | VHFFKNIVT | VHFFKNIVT | VHFFKNIVT | VHFFKNIVT | VHFFKNIVT | VHFFKNIVT | VHFFKNIVT | VHFFKNIVT | VHFFKNIVT | VHFFKNIVT |
| DRB1*15:01 | ENPVVHFFKNIVTPRGGSGGGGG | VHFFKNIVT | VHFFKNIVT | VHFFKNIVT | VHFFKNIVT | VHFFKNIVT | VHFFKNIVT | VHFFKNIVT | VHFFKNIVT | VHFFKNIVT | VHFFKNIVT | VHFFKNIVT | VHFFKNIVT | VHFFKNIVT | VHFFKNIVT |
| DRB5*01:01 | GGVYHFVKKHVHES | YHFVKKHVH | YHFVKKHVH | YHFVKKHVH | YHFVKKHVH | YHFVKKHVH | YHFVKKHVH | YHFVKKHVH | YHFVKKHVH | YHFVKKHVH | YHFVKKHVH | YHFVKKHVH | YHFVKKHVH | FVKKHVHES | YHFVKKHVH |
| DRB5*01:01 | NPVVHFFKNIVTPRTPPPSQ | FKNIVTPRT | FKNIVTPRT | FKNIVTPRT | FKNIVTPRT | FKNIVTPRT | FKNIVTPRT | FKNIVTPRT | FKNIVTPRT | FKNIVTPRT | FKNIVTPRT | FKNIVTPRT | FKNIVTPRT | FKNIVTPRT | FKNIVTPRT |
| DRB5*01:01 | VHFFKNIVTPRTPGG | FKNIVTPRT | FKNIVTPRT | FKNIVTPRT | FKNIVTPRT | FKNIVTPRT | FKNIVTPRT | FKNIVTPRT | FKNIVTPRT | FKNIVTPRT | FKNIVTPRT | FKNIVTPRT | FFKNIVTPR | FKNIVTPRT | FKNIVTPRT |
| Matching | | | 87% | 87% | 93% | 93% | 93% | 93% | 93% | 93% | 93% | 93% | 93% | 87% | 93% |

**Supplementary Figure S9. Binding core predictions from similarity-based fingerprints compared to experimental references.** Predicted versus reference binding cores for 15 HLA-DR restricted peptides as determined from protein structures (Protein Data Bank, retrieved from Ref. (14)). Columns indicate HLA-DR restriction, peptide sequence, reference core (experimentally determined), and predicted cores for each peptide-level fingerprint. Green: matching predictions; red: divergent predictions.

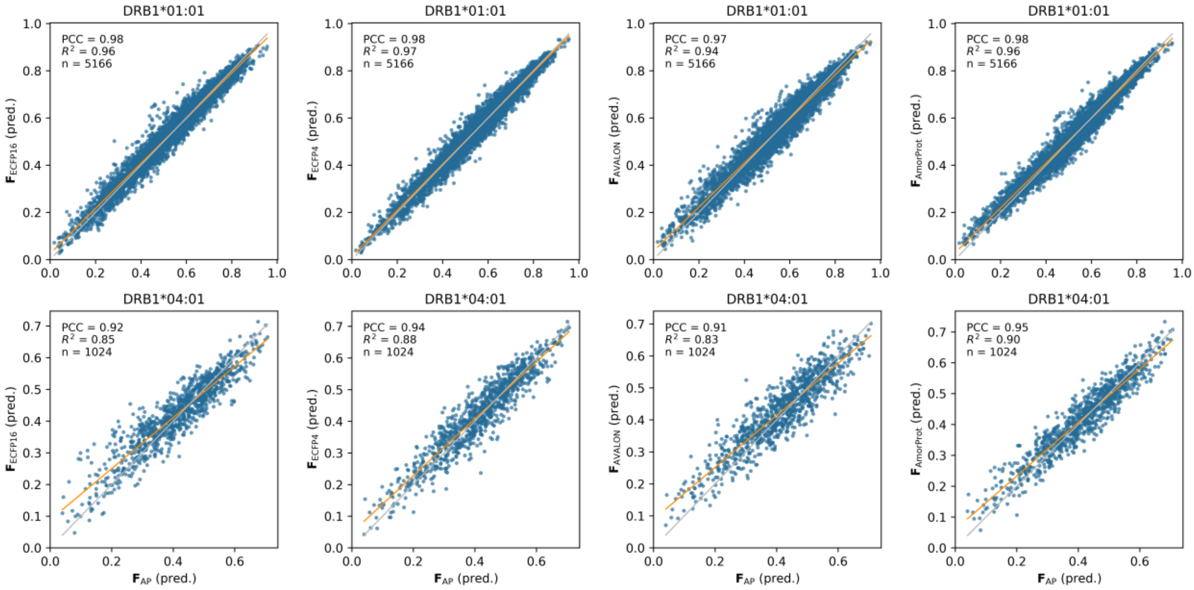


**Supplementary Figure S10. Correlation among $ba_{score}$ predictions of $F_{AP}$ against selected direct-encoding fingerprints for DRB1*01:01 and DRB1*04:01.** Blue dots: natural peptides; yellow line: fitted regression; gray line: identity line. Each panel shows results PCC and $R^2$ performance metrics.

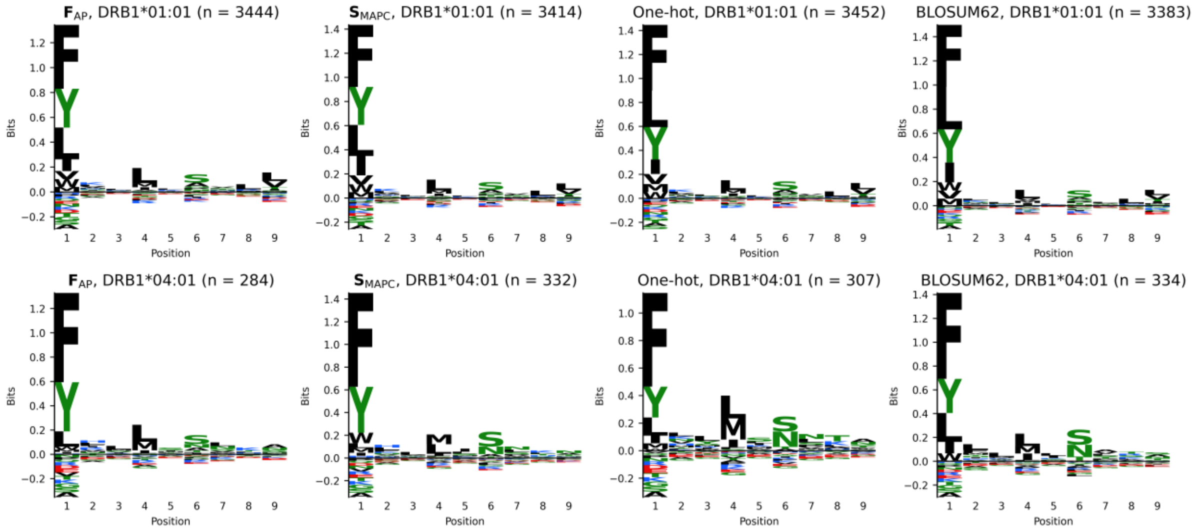


**Supplementary Figure S11. Predicted binding motifs for MHC class II alleles.** Sequence logos generated from out-of-fold predictions for peptides predicted as binders ($ba_{score} \geq 0.5$), considering $\mathbf{F}_{AP}$, $\mathbf{S}_{MAPC}$, one hot and BLOSUM62 descriptors. Logos display amino acid frequencies at each position, with letter height proportional to information content.

### 3. Comparison with TEPITOPE-derived methods

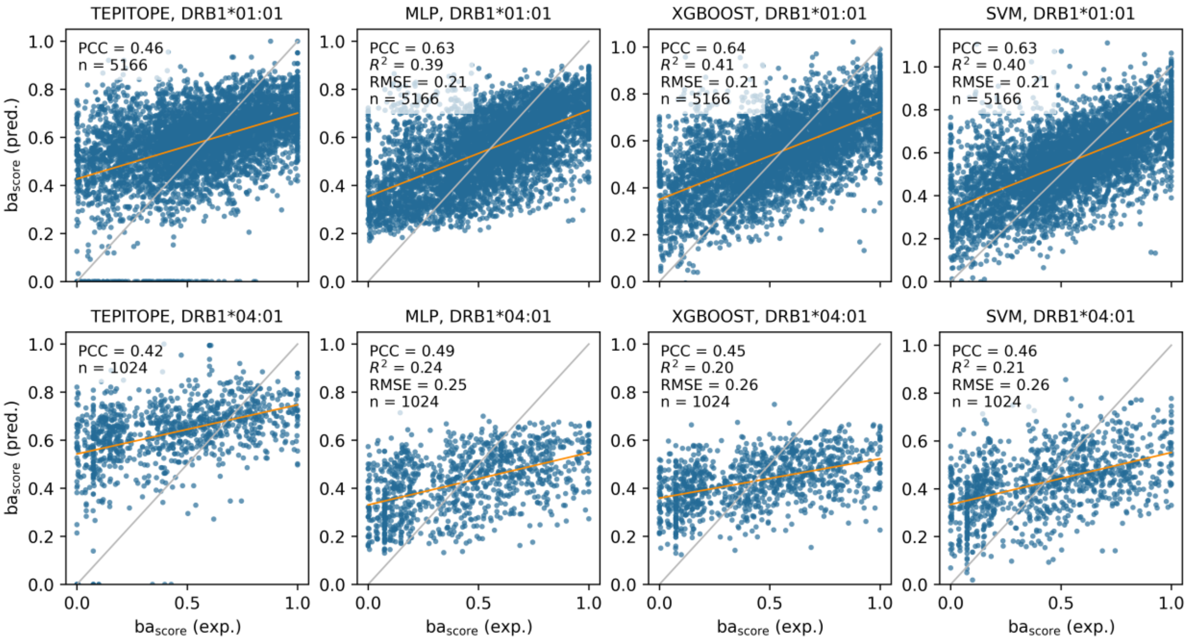


**Supplementary Figure S12. Prediction performances for TEPITOPE-derived methods.** Out-of-fold predictions for DRB1*01:01 and DRB1*04:01 using TEPITOPE-based pocket profile methods. Blue dots represent natural peptides. Yellow line: fitted regression; gray line: identity (perfect prediction). PCC values quantify ranking accuracy. TEPITOPE predictions were min-max scaled to [0,1] range for visual comparability with $ba_{score}$ values. For raw TEPITOPE predictions, $R^2$ values are not reported due to scale differences between TEPITOPE scores and $ba_{score}$ measurements. MLP, XGBoost and SVM models were trained in cross-validation, following the same methodological strategy described in the manuscript. Hyperparameters were optimized using Optuna, as reported in **Supplementary Table S1**.

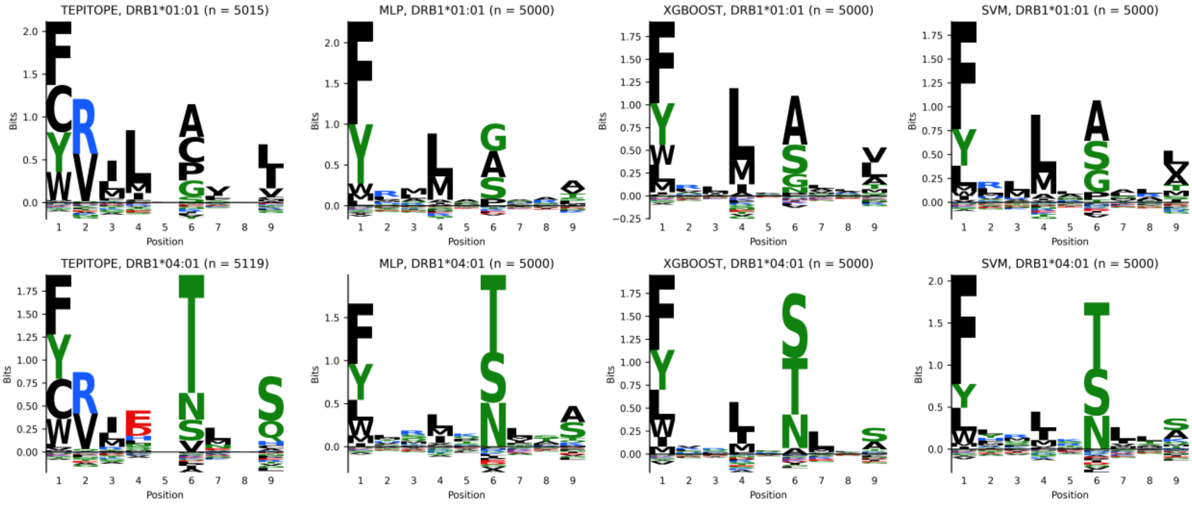


**Supplementary Figure S13. Predicted binding motifs for TEPITOPE-derived methods.** Sequence logos generated from the top 1% predicted binders among randomly sampled peptides from the UniProt human proteome for selected MHC class II alleles. Logos display amino acid frequencies at each position, with letter height proportional to information content.

**Supplementary Table S1. Hyperparameter search space for Optuna optimization of MLP, XGBoost and SVM models.** MLP was implemented using PyTorch; XGBoost and SVM were implemented using the XGBoost Python package and scikit-learn, respectively. Hyperparameters were optimized with Optuna (20 trials), considering pruning and minimizing MSE on cross-validation sets. Early stopping was applied for XGBoost and MLP models.

| Model | Hyperparameter | Search Space |
|---|---|---|
| **MLP** | hidden layer dimension | [8, 16, 32, 64] (categorical) |
| | dropout rate | [0.3, 0.7] (uniform float) |
| | learning rate | [$10^{-4}$, $10^{-2}$] (log-uniform float) |
| | weight decay | [$10^{-5}$, $10^{-2}$] (log-uniform float) |
| **XGBoost** | n_estimators | [50, 300] with step 25 (integer) |
| | max_depth | [2, 6] (integer) |
| | min_child_weight | [3, 10] (integer) |
| | learning_rate | [$10^{-2}$, $10^{-1}$] (log-uniform float) |
| | subsample | [0.5, 0.9] (uniform float) |
| | colsample_bytree | [0.5, 0.9] (uniform float) |
| | reg_alpha (L1) | [0.0, 10.0] (uniform float) |
| | reg_lambda (L2) | [1.0, 10.0] (uniform float) |
| **SVM** | C | [$10^{-1}$, $10^{1}$] (log-uniform float) |
| | epsilon | [$10^{-3}$, $5\times10^{-1}$] (log-uniform float) |
| | kernel | [rbf, linear] (categorical) |
| | gamma | [scale, auto] (categorical), optimized only when kernel = 'rbf' |

### 4. Supplementary results for direct-encoding and similarity-based fingerprints (pMHCI)

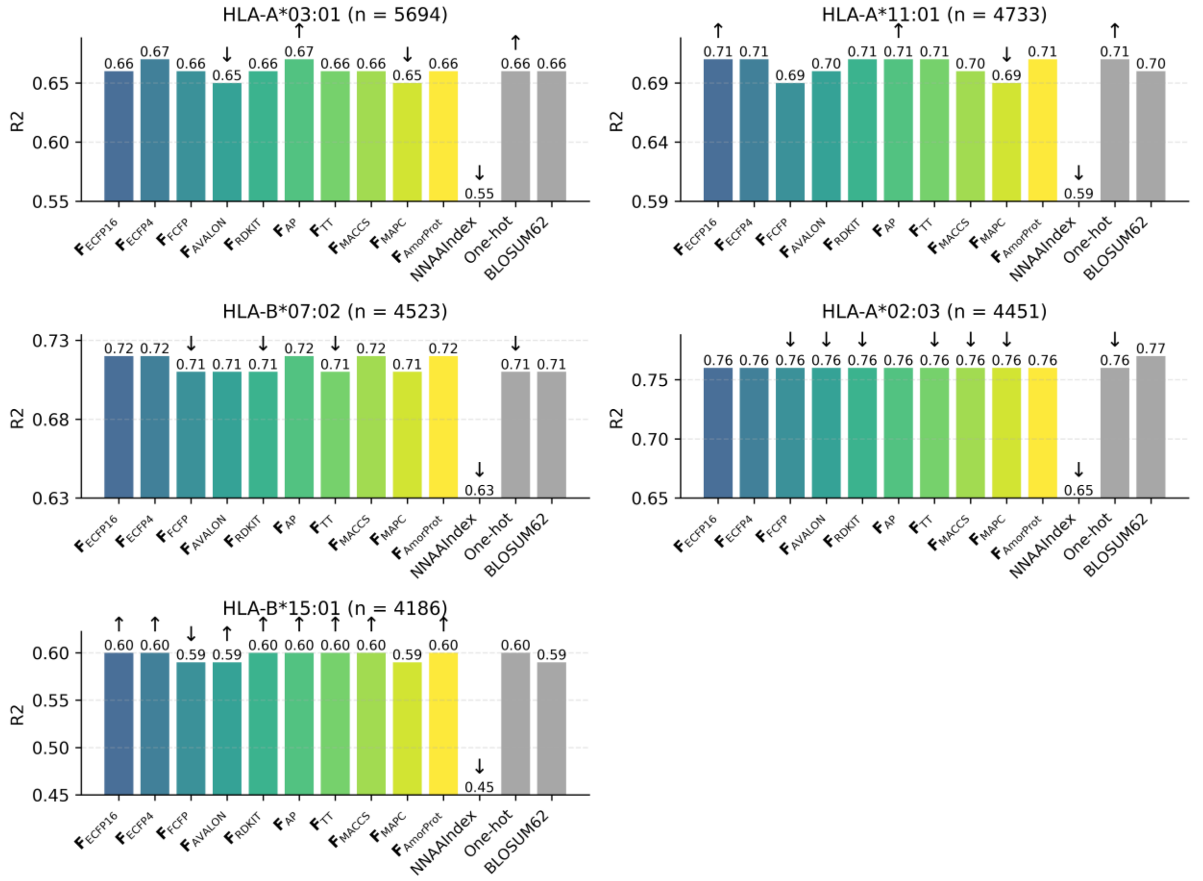


**Supplementary Figure S14. $R^2$ performance of direct-encoding fingerprints for MHC class I binding prediction.** $R^2$ values from out-of-fold predictions for natural peptides binding to selected HLA-A and HLA-B alleles. Each subplot shows results for one allele with sample size indicated in the title. Gray bars represent sequence-based descriptors; coloured bars represent direct-encoding fingerprints. Within each subplot, $R^2$ values are displayed relative to the lowest-performing descriptor to highlight performance differences. Upward arrows indicate significantly better performance than BLOSUM62; downward arrows indicate significantly worse performance than BLOSUM62; absence of arrows indicates no significant difference from BLOSUM62 (paired Wilcoxon tests on per-peptide squared errors, FDR < 0.05, Benjamini-Hochberg correction for 12 pairwise comparisons per allele).

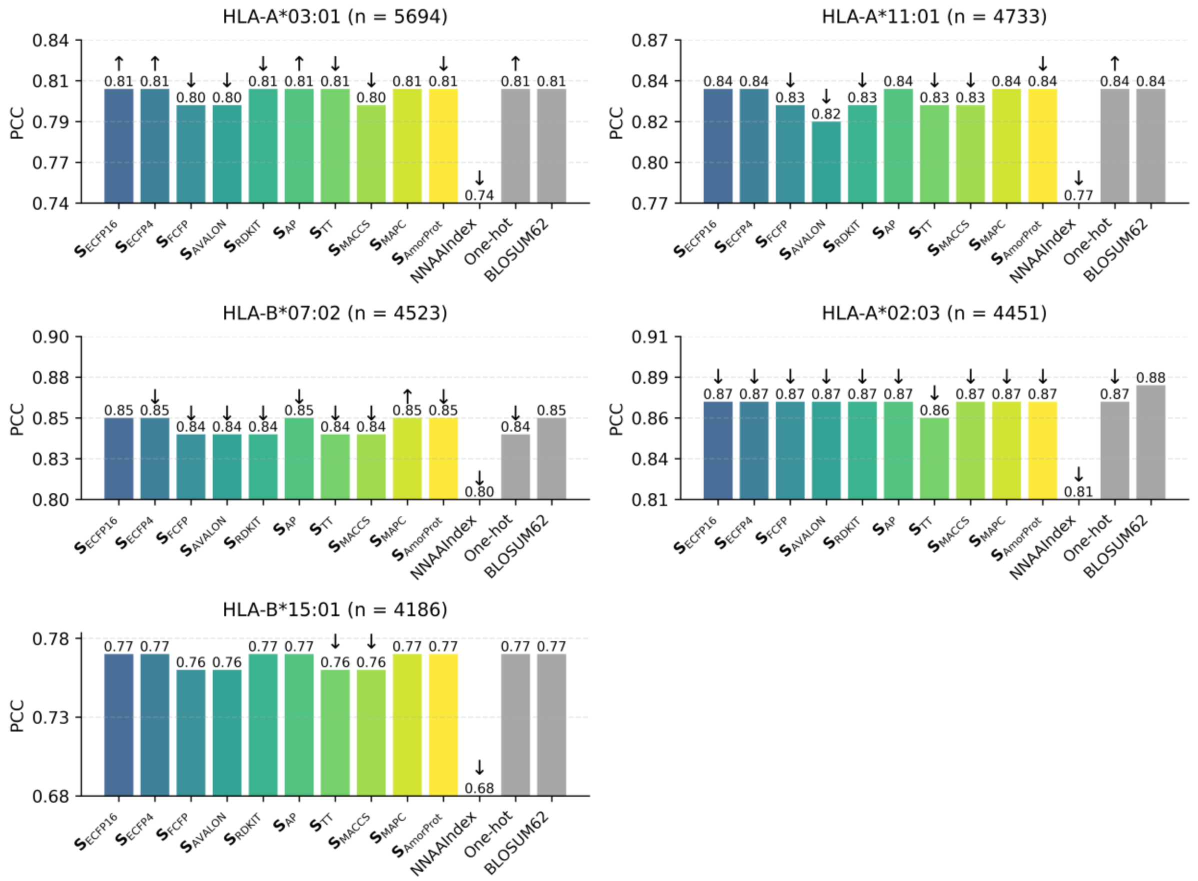


**Supplementary Figure S15. PCC performance of similarity-based fingerprints for MHC class I binding prediction.** PCC values from out-of-fold predictions for natural peptides binding to selected HLA-A and HLA-B alleles. Each subplot shows results for one allele with sample size indicated in the title. Gray bars represent sequence-based descriptors; coloured bars represent similarity-based fingerprints. Within each subplot, PCC values are displayed relative to the lowest-performing descriptor to highlight performance differences. Upward arrows indicate significantly better performance than BLOSUM62; downward arrows indicate significantly worse performance than BLOSUM62; absence of arrows indicates no significant difference from BLOSUM62 (paired Wilcoxon tests on per-peptide squared errors, FDR < 0.05, Benjamini-Hochberg correction for 12 pairwise comparisons per allele).

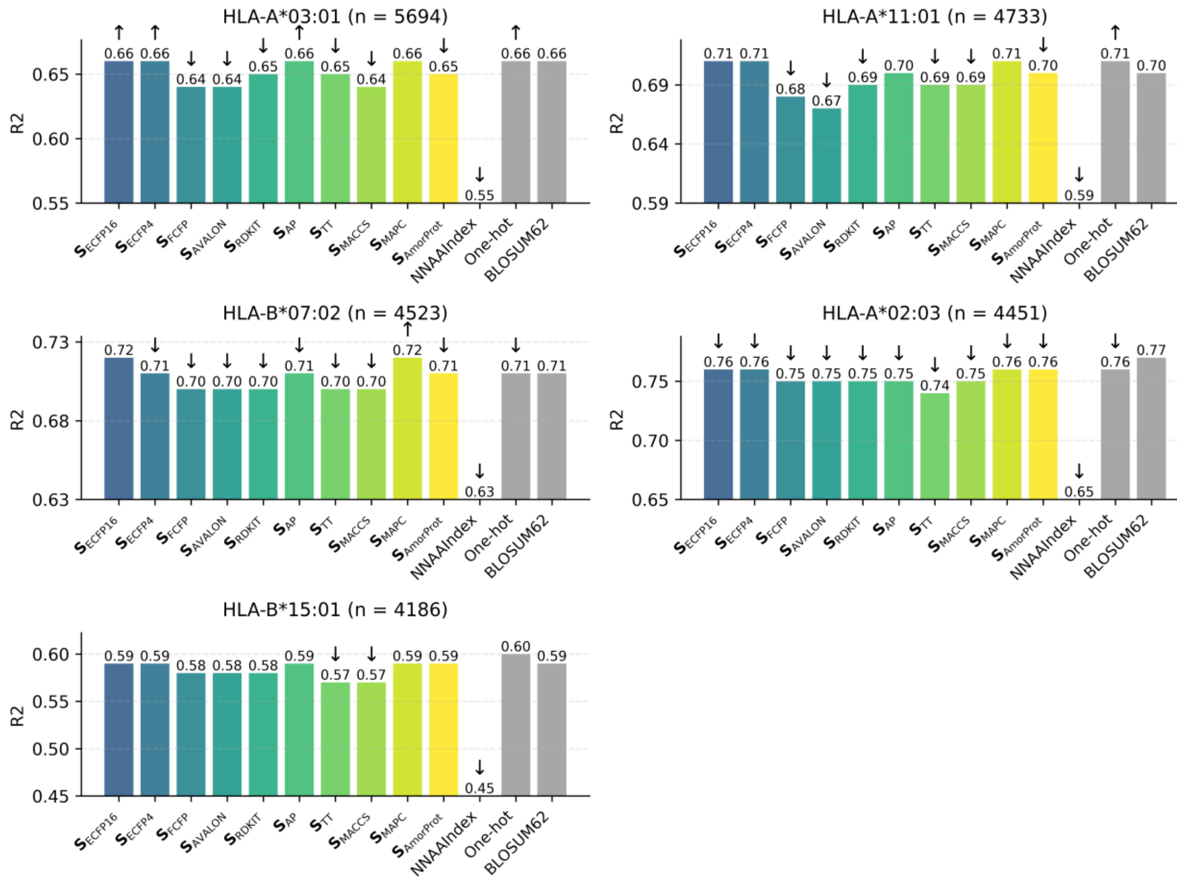


**Supplementary Figure S16. $R^2$ performance of similarity-based fingerprints for MHC class I binding prediction.** $R^2$ values from out-of-fold predictions for natural peptides binding to selected HLA-A and HLA-B alleles. Each subplot shows results for one allele with sample size indicated in the title. Gray bars represent sequence-based descriptors; coloured bars represent similarity-based fingerprints. Within each subplot, $R^2$ values are displayed relative to the lowest-performing descriptor to highlight performance differences. Upward arrows indicate significantly better performance than BLOSUM62; downward arrows indicate significantly worse performance than BLOSUM62; absence of arrows indicates no significant difference from BLOSUM62 (paired Wilcoxon tests on per-peptide squared errors, FDR < 0.05, Benjamini-Hochberg correction for 12 pairwise comparisons per allele).

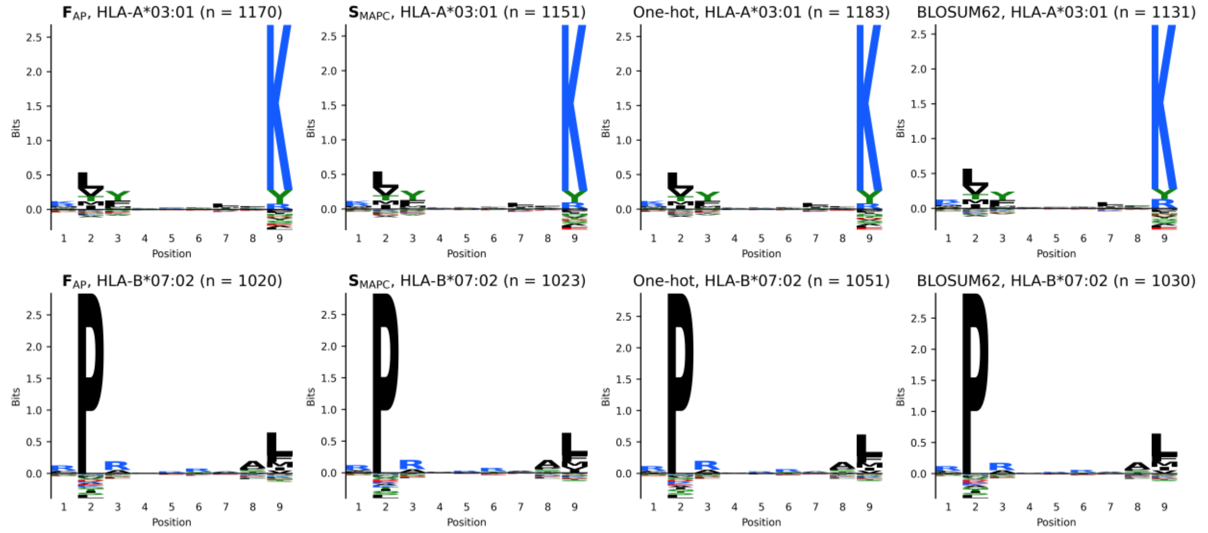


**Supplementary Figure S17. Predicted binding motifs for MHC class I alleles.** Sequence logos generated from out-of-fold predictions for peptides predicted as binders ($ba_{score} \geq 0.5$), considering $\mathbf{F}_{AP}$, $\mathbf{S}_{MAPC}$, one hot and BLOSUM62 descriptors. Logos display amino acid frequencies at each position, with letter height proportional to information content.

### 5. Evaluating the importance of including chemical information

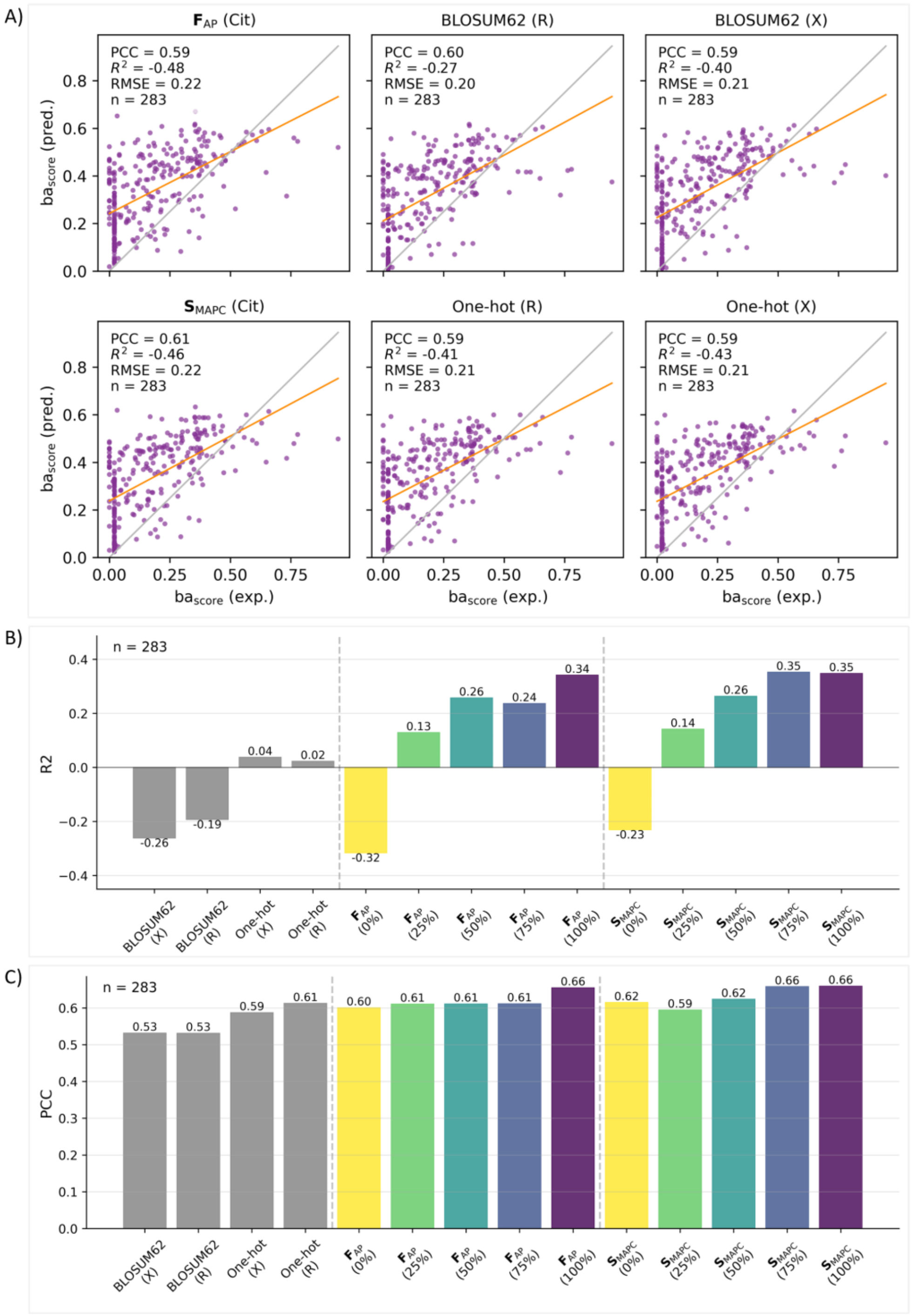


**Supplementary Figure S18. Impact of training data composition on prediction performance for citrullinated peptides (DRB1*04:01).** (**A**) Prediction performance on citrullinated peptides using models trained exclusively on canonical peptides. Yellow line: fitted regression; gray line: identity (perfect prediction). Four descriptors are compared: $\mathbf{F}_{AP}$ and $\mathbf{S}_{MAPC}$ encode citrulline chemically via fingerprints; one-hot and BLOSUM62 encode citrulline as 'X' (zero vector) or 'R' (arginine). (**B**) $R^2$ and (**C**) PCC as a function of citrullinated peptide proportion in training data (0%, 25%, 50%, 75%, 100%). Models were evaluated using 5-fold cross-validation. Gray bars: BLOSUM62 and one-hot encodings trained only on canonical peptides, with citrulline encoded as 'X' or 'R'; coloured bars: $\mathbf{F}_{AP}$ and $\mathbf{S}_{MAPC}$ performance at each training composition.

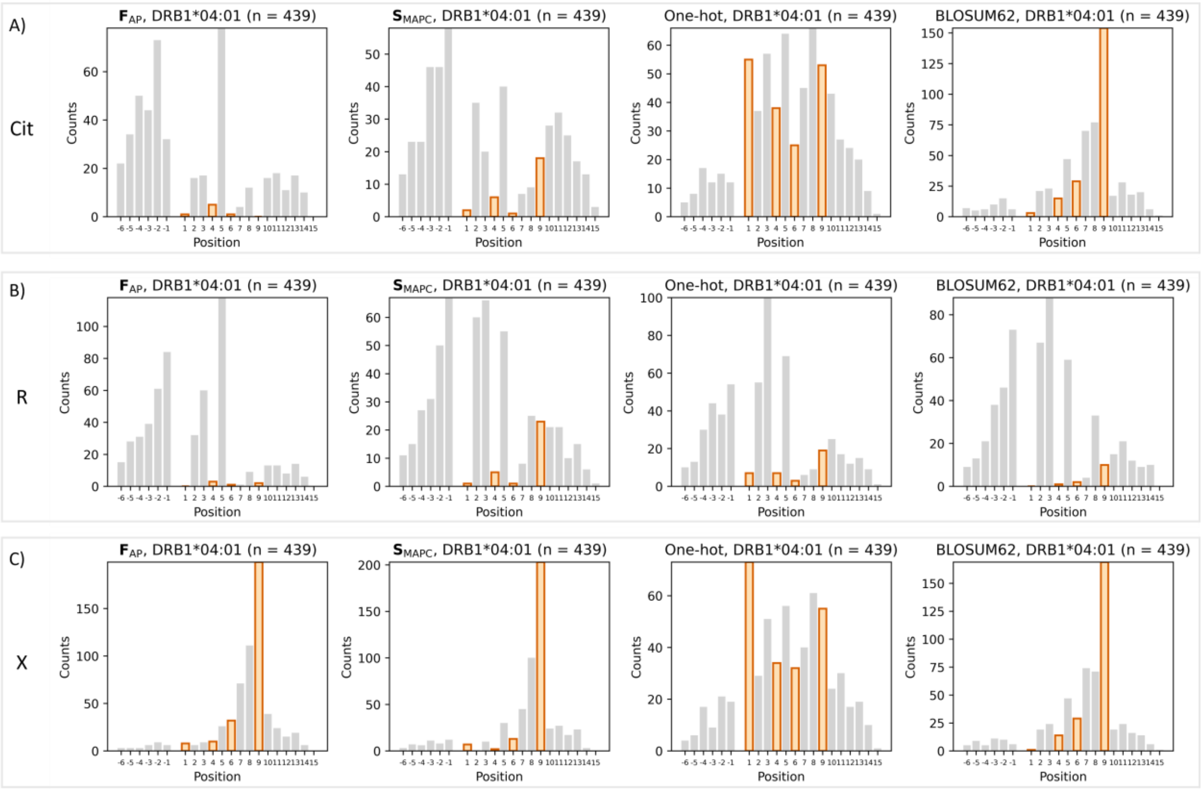


**Supplementary Figure S19. Citrulline positional frequency in predicted binders for DRB1*04:01 under different encoding strategies.** Positional occurrence of citrulline in top 1% predicted binders (n = 439) from citrullinated peptides (Rebak et al. (26)), comparing predictions from models trained with citrulline encoded as: (**A**) citrulline (chemical fingerprint for $\mathbf{F}_{AP}$ and $\mathbf{S}_{MAPC}$ and zero-vector for one-hot and BLOSUM62), (**B**) arginine (natural analogue), or (**C**) 'X' (unknown residue). X-axis shows position relative to the 9-mer binding core: positions 1-9 indicate core positions, negative numbers indicate N-terminal flanking positions, and numbers >9 indicate C-terminal flanking positions. Orange bars highlight frequency at anchor positions (P1, P4, P6, P9); gray bars show frequency at all other positions.
